# Machine learning-assisted directed evolution of plant Rubisco

**DOI:** 10.64898/2026.07.25.740226

**Authors:** Julie L. McDonald, Jiacheng Lin, Yunlong Zhao, Brian L. Hie, Rosemary Birch, Mary Gehring, Bryan D. Bryson, Spencer M. Whitney, Matthew D. Shoulders, Robert H. Wilson

## Abstract

Ribulose-1,5-bisphosphate carboxylase/oxygenase (Rubisco) is foundational to life on Earth, catalyzing carbon dioxide (CO_2_) fixation to generate biomass. However, Rubisco is a slow and inefficient enzyme that has proven challenging to engineer. We applied the structure-informed machine learning (ML) model ESM-IF1 to identify plausible amino acid sites in the large subunit of *Nicotiana tabacum* Rubisco to target for directed evolution. ML-assisted library design followed by selection in Rubisco-dependent *Escherichia coli* identified multiple enriched variants displaying improved catalytic efficiency. Several improved variants carried amino acid changes not found in the evolutionary lineage of plants, despite being assembly competent in plant chloroplasts, demonstrating that ML-assisted protein design can explore functional sequence space beyond what is observed from natural sequence diversity. Most prominently, the T391I substitution improved carboxylation rate by 29% and aerobic carboxylation efficiency by 43%. Our findings demonstrate the utility of ML-assisted evolution for engineering Rubisco with improved carboxylation efficiency and potential for enhancing crop productivity.

## INTRODUCTION

Rubisco is the primary CO_2_-fixing enzyme on Earth, yet it has several shortcomings that limit carbon acquisition, particularly in land plants (*1-3*). Rubisco carboxylates its five-carbon substrate ribulose-1,5-bisphosphate (RuBP) slowly, necessitating its excessive production in the leaves of plants where Rubisco comprises 20–50% of soluble protein (*4*). Further, Rubisco catalyzes RuBP oxygenation, producing 2-phosphoglycolate that C_3_ plants metabolize through photorespiration at a loss of energy, and carbon as CO_2_ (*5*). Rubisco’s catalytic limitations constrain photosynthesis in major C_3_ crops such as soybean, rice, and wheat, making the enzyme a prominent target for improving productivity (*1*).

Expanding knowledge of Rubisco’s kinetic diversity across plant and non-plant lineages has reshaped views of its adaptive potential and called into question the long-held “catalytic trade-off”, where gains in 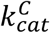 were thought to only arise at the expense of reducing CO_2_/O_2_ specificity (*S*_C/O_) (*6*). Evidence now indicates that this apparent trade-off within vascular plant isoforms is weaker than initially expected (*6*) and may largely reflect the enzyme’s slow natural evolutionary pace and historical adaptation to its CO_2_ environment (*7-10*). Laboratory engineering, freed from these evolutionary constraints, has been employed to push different Rubisco lineages beyond their natural limits, yielding amino acid substitutions that enhance carboxylation kinetics (*11-15*). Among these engineered variants are single substitutions that substantially increase 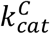 without compromising, and sometimes enhancing, *S*_C/O_ – thus demonstrating that Rubisco’s catalytic landscape is more flexible than previously hypothesized (*16*). The biophysical limits of Rubisco’s carboxylase and oxygenase activities remain a key topic of research relevant to understanding the enzyme’s adaptation to the emergence of atmospheric oxygen and independent evolution of CO_2_ concentrating mechanisms in diverse branches of photosynthetic life (*17, 18*) and reduced inhibition by oxygen observed within the red lineage of Form ID Rubisco (*19*).

Engineering of plant Rubisco was historically bottlenecked by an inability to express the enzyme recombinantly in a tractable laboratory host, owing to a dependence on multiple specialized biogenesis chaperones (*20, 21*). Co-expression of these chaperones alongside the Rubisco large (RbcL) and small (RbcS) subunits permits holoenzyme production in *Escherichia coli* for several dicot (*21-25*), monocot (*26*), and non-vascular basal plants (*27*).

Expression in *E. coli* of ancestral *Solanaceae* Rubisco reconstructions yielded enzymes with improved carboxylation efficiency relative to *Nicotiana tabacum* (tobacco) Rubisco, mediated by multiple substitutions in RbcL and RbcS (*22*). More recently, directed evolution of tobacco Rubisco using random mutagenesis and Rubisco-dependent *E. coli* (RDE) screening was successful in identifying a single *Nt*RbcL residue change (M116L) that increased 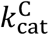 and *S*_C/O_ (*12*). Although the M116L substitution did not improve carboxylation efficiency, the findings show that high-throughput engineering of plant Rubisco is possible, and that directed evolution may eventually offer catalytic improvements to plant Rubisco similar to those already achieved with cyanobacterial (*14*), proteobacterial (*11, 28*) and archaeal (*13*) isoforms.

Traditional directed evolution approaches face a vast sequence space that is impossible to search completely through random mutagenesis. For more efficient library design strategies, protein engineering campaigns have begun to turn to protein language models (pLMs). Such models are pre-trained on vast libraries of sequence and structure information, leading to models that can deliver improved protein functions, ranging from protein–protein interactions to catalytic enhancement (*29-39*). Related to Rubisco in particular, a supervised Gaussian process model was recently applied to predict 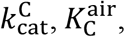, and *S*_C/O_ of many natural Rubisco sequences (*40*), supporting the specific application of machine learning (ML) for Rubisco engineering. However, whether pLMs can identify Rubisco substitutions that improve catalysis remains unclear.

Here, we apply pre-trained, unsupervised ML models to recommend single amino acid substitutions in tobacco Rubisco with high evolutionary and structural likelihood. Though this likelihood score is a term that is only representative of evolutionary plausibility, improvements in likelihood have been observed to translate to increased substrate affinity, kinetics, catalytic activity, and thermostability (*34, 35*), making it a suitable parameter to optimize during Rubisco evolution. We use log-likelihood scores to inform construction of a site-saturated *Nt*RbcL variant library, which proved markedly more efficient than random mutagenesis and enabled the recovery of multiple variants with higher carboxylation efficiency using an RDE selection system (*14*). Several enriched variants incorporated substitutions absent from known vascular plant Rubisco enzymes, underscoring the ability of ML-assisted evolution to explore rare sequence solutions.

These substitutions included the most successful variant, *Nt*RbcL^T391I^, which displayed a 29% faster 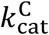 and 43% improvement in aerobic carboxylation efficiency 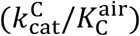. Using a transgenic *Nicotiana benthamiana* line dependent on bacterial *Rhodospirillum rubrum* Rubisco (RbcM) for growth, we further confirmed that ML-recommended, catalytically improved *Nt*RbcL variants could assemble correctly in chloroplasts. Collectively, these results underscore the potential of ML-assisted Rubisco engineering to improve CO_2_ fixation catalysis and, if translated to stably transformed vascular plants, to strengthen crop photosynthesis and productivity.

## RESULTS

### Structure-informed *in silico* deep mutational scan of *Nt*RbcL

We first tested whether the sequence-only model ensemble ESM-1bv (*41, 42*), which has previously been used to inform directed evolution of other proteins (*34*), could effectively recommend beneficial substitutions in the catalytic large subunit of tobacco Rubisco (table S1). ESM-1bv generally suggested amino acid substitutions already common in vascular plant Rubisco outside *Solanaceae* (fig. S1). However, such phylogenetically widespread substitutions are not reliable indicators of kinetic improvement, given that carboxylation efficiency shows little correlation with evolutionary relatedness. That is, highly diverged land plant lineages often display near identical kinetic profiles (*43*). Furthermore, physiological (*44*) and environmental adaptation (*45*) can generate substantial kinetic variation even among closely related species. These considerations suggested to us that a sequence-only model may be insufficient to identify mutations that meaningfully enhance *Nt*RbcL catalysis.

To ground recommendations within the overall architecture and catalytic mechanism of Rubisco, we moved beyond sequence-only models to explore a protein structure-informed ML-based design strategy. ESM-IF1 (*35, 46*), trained on 12 million predicted structures generated by AlphaFold2 (*47*), is designed to generate a protein’s amino acid sequence from an input structural backbone (*48*), meaning that it attempts to predict a set of optimized sequences best adapted to conserved folds. We used ESM-IF1 to score the log-likelihood of single amino acid substitutions at each position in *Nt*RbcL relative to wild-type (**Fig. 1A** and table S1) using crystal structures of tobacco Rubisco in both the unactivated ‘open’ (PDBID: 1RLC) (*49*) and activated ‘closed’ (PDBID: 4RUB) (*50*) configurations. The closed structure contains additional backbone structural information in *Nt*RbcL due to ordering of the C-terminus over the catalytic Loop 6 when Rubisco is activated and substrate bound (*51*). Rubisco in both structures is bound to 2-carboxyarabinitol-1,5-bisphosphate (CABP), a non-hydrolysable analog of the carboxylation transition state. We reasoned that supplying bound structures to ESM-IF1 would help to avoid recommendations that alter the backbone conformation in ways that preclude substrate binding (**Fig. 1B**).

**Fig. 1.**
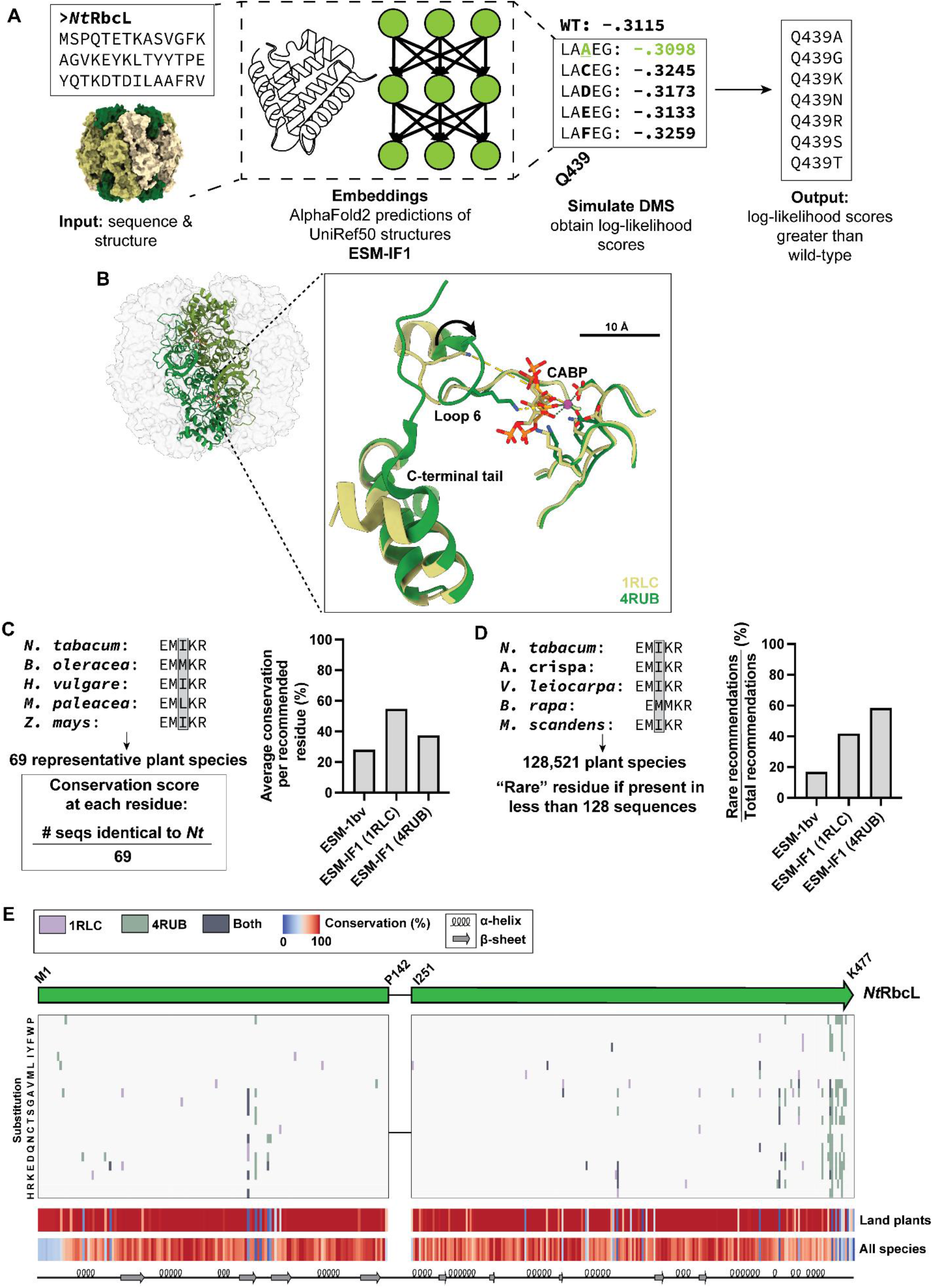
Evolution of *Nt*RbcL using ESM-IF1 *in silico* deep mutational scanning (DMS). (**A**) ESM-IF1 inputs structural information alongside protein sequence. An *in silico* deep mutational scan resulted in several recommended variants with log-likelihood scores greater than wild-type *Nt*RbcL. (**B**) Input structures used for ESM-IF1. PDBID 4RUB differs from PDBID 1RLC by activation (carbamylation) of the catalytic lysine, causing closure of Loop 6 towards the bound substrate and compaction of the C-terminus towards the active site. (**C**) Average conservation of amino acid loci recommended by ESM-1bv and ESM-IF1. Full species names and RbcL GenBank accession codes are listed in (table S2). (**D**) Percentage of the top 24 ESM-1bv and ESM-IF1-recommended substitutions that occur rarely (<0.01% of the time) in sequenced plant Rubisco. (**E**) Location of recommended substitutions in the *Nt*RbcL sequence using different input structures. Conservation and secondary structure at each residue is shown. Approximate boundaries of the N-terminal, TIM barrel, and C-terminal domains are indicated. Sequence is not shown from residue 143–250 as no ML recommendations were made in this region.

In total, 202 substitutions at 49 positions across both structures displayed log-likelihood scores greater than the corresponding wild-type amino acid in *Nt*RbcL. Of these substitutions, 29 were recommended using both structural inputs and 20 overlapped with recommendations from the ESM-1bv ensemble (table S1). Recommendations from ESM-IF1 were, on average, located at more highly conserved residues than recommendations from ESM-1bv. We calculated a conservation score for each residue in *Nt*RbcL using a diverse dataset of land plant sequences (*9*), where the score is the percentage of sequences in the dataset that contain the same amino acid at a given position as *Nt*RbcL (**Fig. 1C**). The average conservation score of the 18 amino acid sites that comprise the 24 amino acid substitutions recommended by ESM-1bv was 28%. For consistency with the ESM-1bv predictions, we calculated the average conservation score of sites within the 24 most highly-ranked ESM-IF1 recommendations. These sites had an average conservation score of 37–55% depending on input structure. We also surveyed a larger dataset of approximately 128,000 plant Rubisco sequences to determine how often the models recommended amino acid substitutions that are rarely observed in natural plant Rubisco sequences (**Fig. 1D**). 17% of substitutions recommended by ESM-1bv were found in <0.01% of plant Rubisco sequences, and all rare substitutions were recommended by only one of the six models in the ensemble. In contrast, 42 and 58% of the top 24 ESM-IF1 recommendations were rare for ‘open’ (PDBID: 1RLC) and ‘closed’ (PDBID: 4RUB) structural inputs, respectively.

Many ESM-IF1 recommendations were located in the C-terminus of *Nt*RbcL (**Fig. 1E**), especially for the closed structure. Encouragingly, there were recommendations where congruent substitutions enhance catalytic parameters in distantly related Rubisco isoforms, such as W451F in the archaeal Rubisco from *Methanococcoides burtonii* (as Y471) (*13*). Importantly, no substitutions were predicted with a higher log-likelihood score than wild-type at site F345, a locus that is commonly selected for its solubility-enhancing outcome during directed evolution of related Rubisco from cyanobacteria (*14, 52-54*).

### Preliminary screening and characterization of structure-informed *Nt*RbcL sequence recommendations

We sought to evaluate whether the predicted structure-based substitutions permitted Rubisco folding and function using the RDE2 bacterial screening and selection platform. RDE2 is a second-generation RDE system in which phosphoribulokinase (PRK) is expressed from an arabinose-inducible promoter, causing the accumulation of cytotoxic RuBP that is removed through Rubisco catalysis (*14*). We utilized a two-plasmid system that enables plant Rubisco expression in *E. coli* and is compatible with RDE2 (*12, 24*) to screen tobacco Rubisco variants. As in prior work, the Rubisco plasmid employed, pET-*Nt*LS-Rca, contains a V369A substitution in *Nt*RbcL and is referred to as ‘wild-type’ henceforth. V369A does not substantially impact catalysis, but does improve the enzyme’s solubility in *E. coli* (table S3). The use of an already high-solubility *Nt*RbcL variant helps to ensure selection in RDE2 on the basis of improved kinetic properties rather than merely improved solubility (*11, 14*).

For an initial trial, three substitutions in *Nt*RbcL from 202 recommendations were manually selected. The substitution I465V was chosen because of its previous selection during directed evolution of cyanobacterial Rubisco (*52, 53*), although such selection was potentially due to solubility effects. We also chose two previously uncharacterized charge-switching substitutions, A438R and Q439K, based on their proximity to one another and differences in their natural occurrence. R438 was not observed in natural sequences surveyed and K439 is found only in bacterial Rubisco (**Fig. 2A**), although the similarly positively charged residue R439 is widespread in plant Rubisco.

**Fig. 2.**
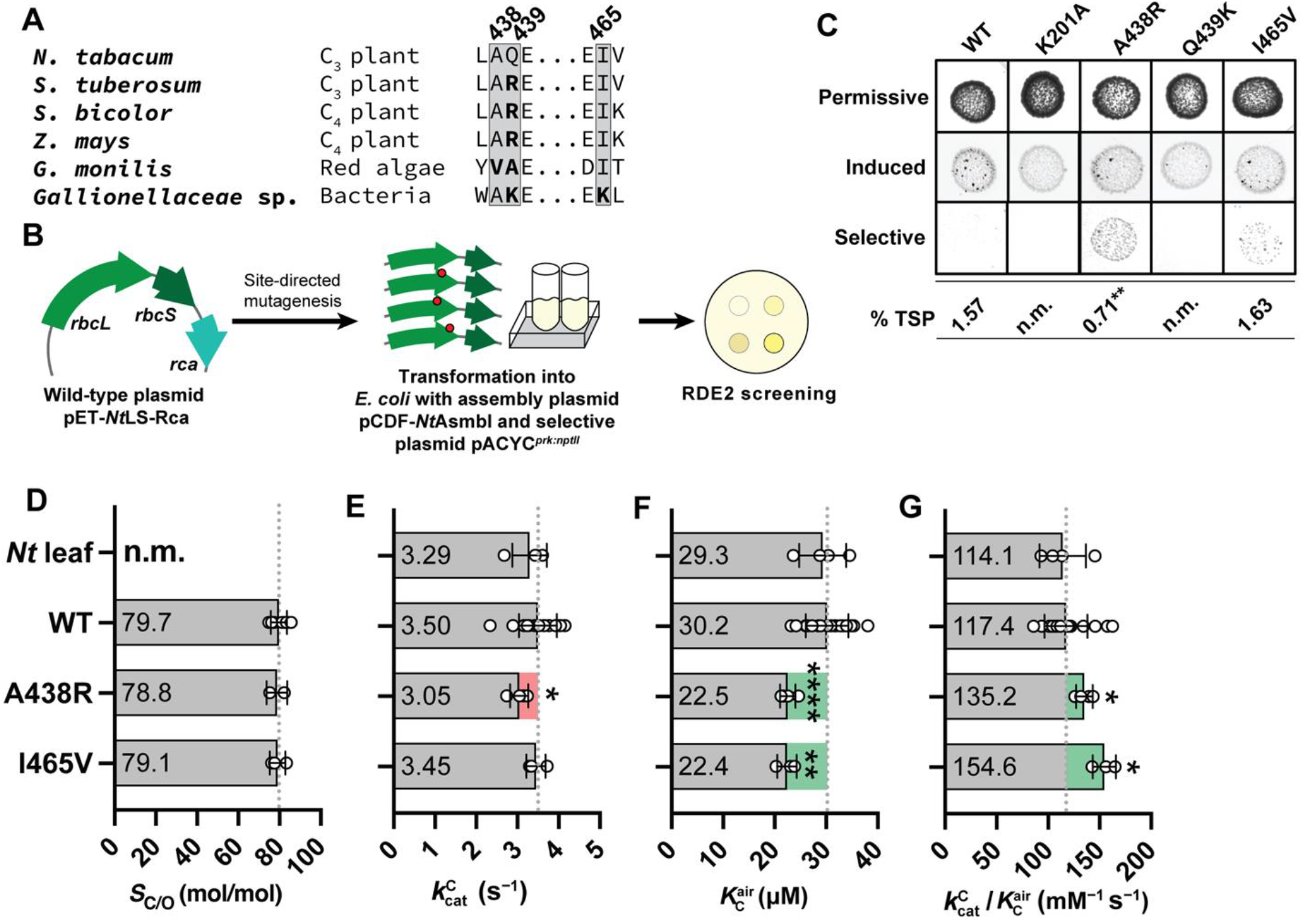
Characterization of select ESM-IF1-recommended NtRbcL variants. (**A**) Sequence alignment of Rubisco across phylogeny. ESM-IF1-recommended sites are boxed in gray and sequence differences relative to *Nt*RbcL are bolded. Full species names and RbcL GenBank accession codes are listed in (table S2). (**B)** Workflow: Plasmids (pET-*Nt*LS-Rca) coding wild-type (WT) or ESM-IF1-recommended point mutations in *Nt*RbcL were transformed into RDE2 containing plasmids pACYC^*prk:nptII*^ for PRK-dependent selection and pCDF-*Nt*Asmbl for assembly of the tobacco Rubisco holoenzyme. (**C)** RDE2 spot tests for comparing Rubisco-facilitated growth on permissive media (no additives), Rubisco-inducing media (0.2 mM IPTG), and Rubisco + PRK-selective media (0.15% [w/v] L-arabinose, 400 µg/mL kanamycin, and 0.2 mM IPTG). Soluble Rubisco content in *E. coli*, quantified by [^14^C]-CABP binding (*55*), is shown as % total soluble protein (% TSP). (**D**) Specificity factor; *S*_C/O_. (**E**) Carboxylation rate; 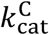. (**F**) Affinity for CO_2_ in air;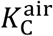. (**G**) Carboxylation efficiency in air; 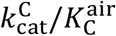. Significance testing for panels **C**–**G** was performed via two-tailed, heteroscedastic *t*-tests of each Rubisco variant relative to wild-type. * = *p* ≤ 0.05; ** = *p* ≤ 0.005; **** = *p* ≤ 0.00005. All error bars represent standard deviation. Replicate data for panels **C**–**G** are listed in (table S4). Dashed lines in panels **D**–**G** represent the mean of wild-type measurements for each parameter.

We generated the *Nt*RbcL A438R, Q439K, and I465V variants using site-directed mutagenesis and screened them using RDE2 alongside an inactive Rubisco negative control (*Nt*RbcL^K201A^; **Fig. 2B**). After four days under RDE selection on strongly selective media containing 0.15% [w/v] L-arabinose in an atmosphere containing 0.5% [v/v] CO_2_ and 21% [v/v] O_2_, *E. coli* expressing the wild-type, *Nt*RbcL^Q439K^, or *Nt*RbcL^K201A^ tobacco Rubisco failed to grow. Cells producing *Nt*RbcL^A438R^ and *Nt*RbcL^I465V^ survived (**Fig. 2C)**. Neither the *Nt*RbcL^A438R^ nor *Nt*RbcL^I465V^ mutants improved Rubisco solubility (**Fig. 2C**), indicating their fitness-enhancing properties were likely catalytic and motivating further characterization of these two variants.

Measurements of 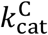 using soluble protein in *E. coli* lysate exhibited high inter-experiment variability relative to *N. tabacum* leaf soluble protein controls (fig. S2). Attempts to purify Rubisco from *E. coli* lysate using a chromatographic purification protocol optimized for Rubisco purification from leaves (*56*) proved low-throughput and resulted in low yields of still heavily contaminated end-product (fig. S3). To simplify the purification process, we adapted a nickel-immobilized metal affinity chromatography (IMAC) purification method previously used to purify *Synechococcus* sp. PCC6301 Rubisco (*52*) by appending an N-terminal cleavable histidine-tagged ubiquitin (^His6^Ubq) to the Rubisco small subunit (*Nt*^His6^UbqRbcS; figure S4A–B). Co-expression of *Nt*RbcL, untagged *Nt*RbcS, and *Nt*^His6^UbqRbcS allowed for the purification of tobacco Rubisco complexes by IMAC prior to cleaving off the ubiquitin tag (fig. S4C) (*57*). Using this method, the isolated Rubisco contained no amino acid epitope tags, was typically >70% pure, and resulted in more reproducible measurements of 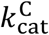 and 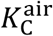 (fig. S2).

Compared with wild-type tobacco Rubisco, the *Nt*RbcL^A438R^ and *Nt*RbcL^I465V^ variants showed no differences in *S*_C/O_ (**Fig. 2D**). The 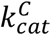 of *Nt*RbcL^A438R^ was slightly decreased relative to wild-type and the 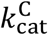 of *Nt*RbcL^I465V^ was unchanged (**Fig. 2E**). However, the apparent CO_2_ affinity 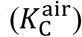 of both *Nt*RbcL^A438R^ and *Nt*RbcL^I465V^ improved by ∼25% (**Fig. 2F**), leading to 15% and 32% improvements in their carboxylation efficiency 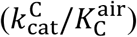, respectively (**Fig. 2G**). Thus, in accordance with previous work screening plant (*12*) and bacterial (*13, 14*) Rubisco variants, improved RDE2 selective growth could be ascribed to an improvement in catalysis.

To more deeply analyze ESM-IF1 recommendations, we generated site-saturated mutagenesis libraries at the 49 recommended *Nt*RbcL amino acid sites (table S1), along with K201 as a negative control. This library of ∼400 variants (**Fig. 3A**) contained at least one substitution at each target site aside from R89, which failed during insert fragment synthesis (table S5). Next-generation sequencing (NGS) coverage of the plasmid was highly variable, especially at *rbc*L (fig. S5).

**Fig. 3.**
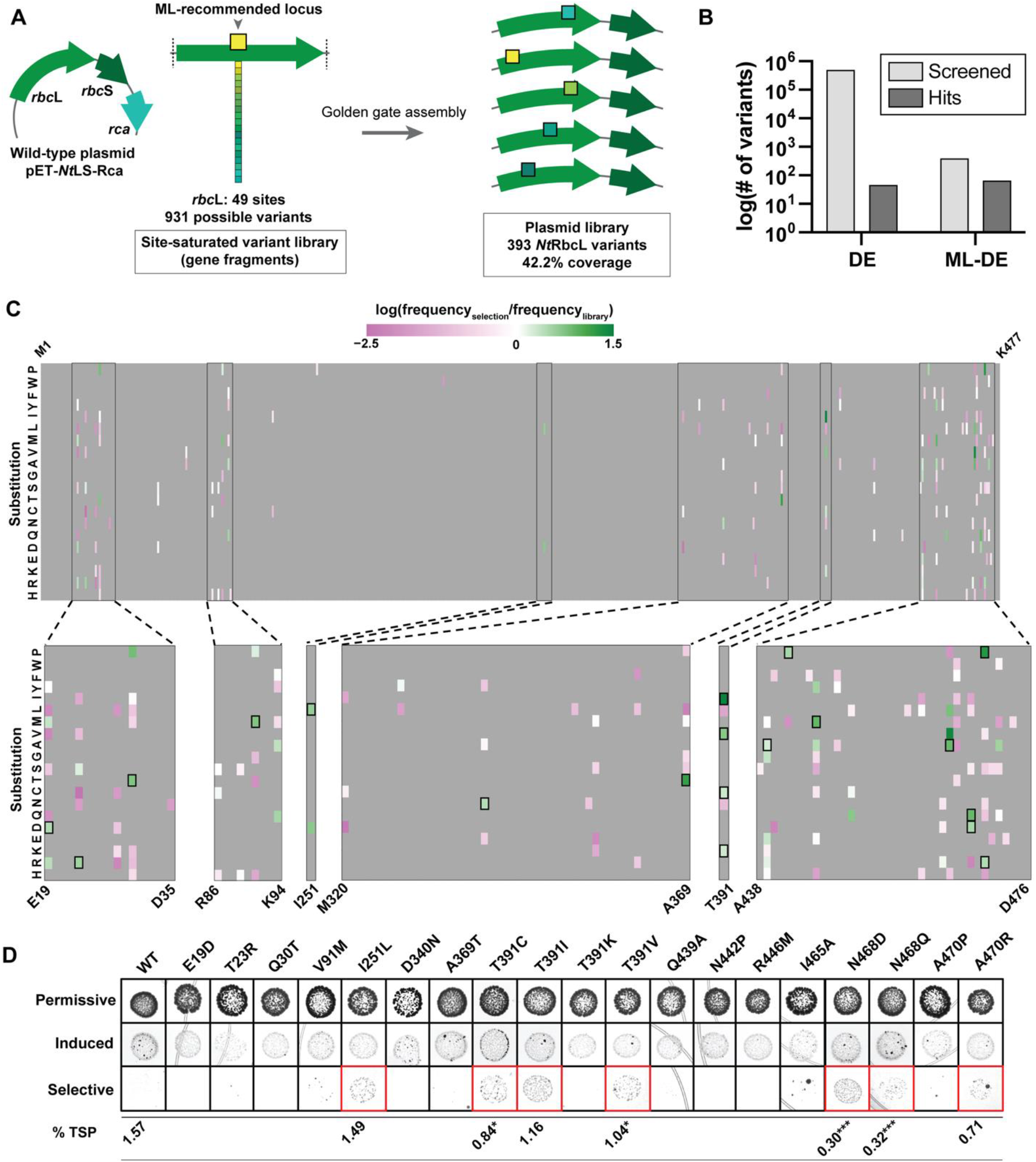
Library screening at ESM-IF1-recommended sites. (**A**) Construction of a site-saturated variant library at 49 sites by Golden Gate assembly. (**B**) Tobacco Rubisco variants screened and isolated as “hits” from ML-assisted directed evolution (ML-DE, this work) and traditional directed evolution (DE; (*12*)). Hits from DE screening were identified as fast-growing colonies in RDE2, while hits from ML-DE screening were identified through enrichment in NGS reads. (**C**) Enrichment of variants in RDE2 selection. Enrichment score was calculated as frequency of the variant on selective media divided by the frequency of the variant in the starting library. Heatmap represents log scale enrichment scores (table S5). (**D**) RDE2 rescreening of enriched variants. Media conditions are equivalent to those in (**Fig. 2C**). Soluble Rubisco content in *E. coli* for each measured variant is shown. Significance is shown relative to wild-type tobacco Rubisco as analyzed via a two-tailed, heteroscedastic *t*-test. * = *p* ≤ 0.05; *** = *p* ≤ 0.0005.

Excision of *rbc*L was apparent during library generation, indicating that cytotoxicity or plasmid instability from the repetitive T7 promoter and terminator sequences (fig S4A) may have prevented construction of all possible *Nt*RbcL variants.

Library selection in RDE2 followed by NGS found 64 substitutions, including I465V, that were enriched following selection relative to their frequency in the starting library, while substitutions at the catalytically critical K201 locus were consistently depleted (**Fig. 3B–C** and table S5). The overall positive enrichment rate was 16%, 1,700-fold higher compared to prior work selecting from error-prone PCR libraries of *N. tabacum* Rubisco that identified 46 enriched variants from a library of ∼500,000 (*12*) (**Fig. 3B**). ML-assisted library preparation is therefore effective for generating small, streamlined Rubisco libraries that successfully navigate the high degree of sequence conservation present in land plant isoforms (**Fig. 1E**).

Of nineteen chosen Rubisco variants enriched under >0.1% arabinose selection (**Fig. 3C**, black boxes), seven showed improved RDE2 growth relative to wild-type that was not attributable to increased Rubisco solubility (**Fig. 3D**, red boxes). We additionally combined the top five, ten, and twenty most highly enriched point mutations to test for additive benefit to RDE2 growth (fig. S6). The quintuple variant *Nt*RbcL resulted in slightly worse growth in RDE2 compared to the most highly enriched variant (*Nt*RbcL^T391I^), suggesting no apparent improvement to *Nt*RbcL function or solubility with the addition of more substitutions. The variants containing 10 and 20 substitutions failed to grow under selective conditions, indicating that these combinations of substitutions impaired Rubisco folding and/or function.

### ML-assisted engineering achieves improved carboxylation efficiency

*Nt*RbcL containing the single substitutions I251L, T391C, T391I, T391V, N468D, N468Q or A470R were purified and carboxylation catalysis was measured under ambient atmospheric O_2_ (21% [v/v], equilibrated as 253 μM dissolved O_2_) to mimic the physiological conditions of a plant chloroplast (*58*) and to obtain 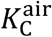 values to model the response of leaf photosynthetic rate (*A*) under varying chloroplast CO_2_ pressures (*C*_c_) (*59*) (**Table 1** and table S4). The carboxylation properties of the most strongly enriched RDE2 variant, *Nt*RbcL^T391I^, were the most improved, with a 29% higher 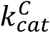 and 11% reduction in 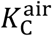 to give a net 43% increase in 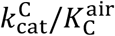 and *Nt*RbcL^T391V^, as well as the previously measured variants *Nt*RbcL^A438R^ and *Nt*RbcL^I465V^ (**Fig. 2G**), also displayed improved 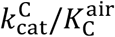 (**Fig. 4A**) owing to reductions to 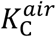 while maintaining a similar 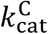 to wild-type. *Nt*RbcL^T391C^, *Nt*RbcL^N468D^, and *Nt*RbcL^N468Q^ displayed a similar catalytic trend of improved 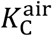, although overall improvement to carboxylation efficiency was not significant for these variants (**Table 1**). Measurements of *S*_C/O_ (**Fig. 4B**) revealed that none of the amino acid changes significantly impacted the overall specificity of the enzyme for CO_2_ over O_2_.

**Table 1.** Kinetic characterization of *Nt*RbcL variants at 25 °C.

| *NtRbcL* | Enrichment score | $k_{\text{cat}}^{\text{C}}$ (s<sup>-1</sup>) | $K_{\text{C}}^{\text{air}}$ (μM) | $k_{\text{cat}}^{\text{C}}/K_{\text{C}}^{\text{air}}$ (mM<sup>-1</sup> s<sup>-1</sup>) | $S_{\text{C/O}}$ |
| --- | --- | --- | --- | --- | --- |
| Wild-type | - | 3.5 ± 0.5 | 30.2 ± 4.1 | 117 ± 21 | 79.7 ± 4.0 |
| I251L | 17.7 | 3.1 ± 0.1* | 22.0 ± 1.2**** | 142 ± 7** | 79.8 ± 3.1 |
| T391C | 8.4 | 3.3 ± 0.3 | 24.9 ± 0.7*** | 131 ± 9 | 79.2 ± 0.6 |
| T391I | 189.3 | 4.5 ± 0.4* | 26.9 ± 1.0* | 168 ± 19* | 79.9 ± 3.1 |
| T391V | 2.2 | 3.3 ± 0.1 | 23.0 ± 1.4*** | 143 ± 12* | 78.7 ± 3.3 |
| A438R | - | 3.0 ± 0.2* | 22.5 ± 1.5**** | 135 ± 8* | 78.8 ± 5.0 |
| I465V | 136.0 | 3.5 ± 0.2 | 22.4 ± 1.9** | 155 ± 11* | 79.1 ± 3.8 |
| N468D | 13.5 | 3.0 ± 0.3 | 20.2 ± 6.4 | 156 ± 32 | 76.7 ± 3.5 |
| N468Q | 36.1 | 3.1 ± 0.3 | 20.7 ± 0.5**** | 152 ± 18 | 80.8 ± 1.3 |
| A470R | 12.5 | 3.6 ± 0.3 | 29.9 ± 4.6 | 124 ± 20 | 77.4 ± 0.9 |

Values provided are means ± standard deviation. Replicate data are provided in (table S4). Statistical significance relative to wild-type was calculated via a two-tailed, heteroscedastic *t*-test. \* = $p \leq 0.05$ ; \*\* = $p \leq 0.005$ ; \*\*\* = $p \leq 0.0005$ ; \*\*\*\* = $p \leq 0.00005$ .

**Fig. 4.**
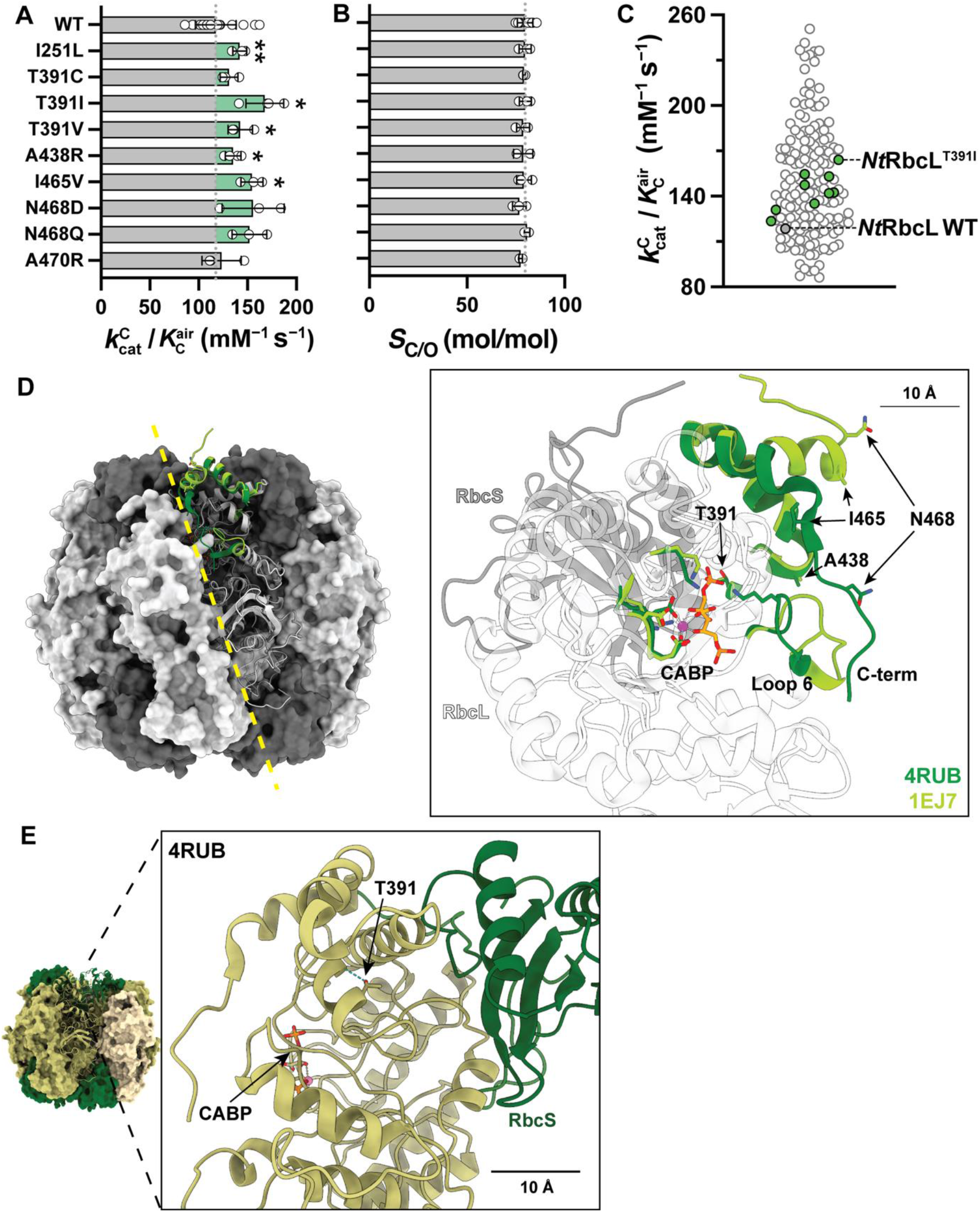
Tobacco Rubisco variants in kinetic and structural contexts. (**A**) Carboxylation efficiency in air 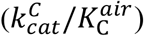 and (**B**) CO_2_/O_2_ specificity factor (*S*_C/O_) for wild-type *Nt*RbcL and variants. Statistical significance relative to wild-type was calculated via a two-tailed, heteroscedastic *t*-test. * = *p* ≤ 0.05; ** = *p* ≤ 0.005. (**C**) Carboxylation efficiencies in air of C_3_ plants, displayed with data collected in this study and from (*60*). *Nt*RbcL variants measured in this work are shown as filled green circles while wild-type *Nt*RbcL is shown as a filled gray circle. (**D**) Structural analysis of amino acids near the *Nt*RbcL active site that enhance carboxylation catalysis. PDBID: 4RUB (*50*) (activated, closed, and CABP-bound) and 1EJ7 (*61*) (unactivated, open, and phosphate-bound). (**E**) Structural contextualization of *Nt*RbcL^T391I^ between *Nt*RbcL and *Nt*RbcS within the Rubisco hexadecamer. PDBID: 4RUB.

Previously analyzed plant Rubisco, especially those from C_3_ plants, occupy a narrow range (*60*) of carboxylation efficiency (86–251 mM^−1^ s^−1^) in their natural aerobic environment (i.e., under 21% [v/v] ambient O_2_ (*9, 60, 62*)). Compared with measured 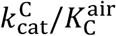 data from previous studies (*60*), we found our ML-assisted directed evolution approach enabled tobacco Rubisco to traverse across 30% of this C_3_ plant Rubisco catalytic landscape via single substitutions alone (**Fig. 4C**). As each naturally evolved amino acid substitution that occurs in RbcL is proposed to increase the enzyme’s anaerobic carboxylation efficiency by only ∼1% (*9*), the ML-based approach taken here identified functional mutations that appear to sidestep this natural constraint (for aerobic carboxylation efficiency).

Given that these beneficial substitution sites were predicted using structural information alongside protein sequence, we aimed to understand the potential impact of these substitutions on the overall structure of the holoenzyme. Several of the catalytically improved variants, specifically A438R, I465V, N468D, and N468Q, are located in the C-terminal portion of the enzyme (**Fig. 4D**), a highly mobile region directly involved in the catalytic mechanism, where it operates to latch Loop 6 in the closed state following RuBP binding. This process requires the breakage and formation of several hydrogen bonds, thereby contributing to the overall energetics of the reaction (*51, 63*). I465 and N468 in particular are positioned at the terminus of the final α-helix of *Nt*RbcL, which becomes more disordered during C-terminal-tail repositioning. I465 contributes to a spiral channel along with residues F467 and F469, which flank N468. This channel is proposed to funnel gas substrates to the active site (*64*) and may explain the observed improvements in 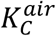 for *Nt*RbcL^I465V^, and by *Nt*RbcL^N468D/Q^, through stabilization or favorable positioning of the F467 and F469 side chains within the funnel. We note that the gas funnel is not an experimentally substantiated feature of Rubisco catalysis and that both F467 and F469 form critical interactions with the assembly chaperone RbcX (*65*) that may alternatively explain their high sequence conservation, and the detrimental influence of N468 mutants on Rubisco solubility (**Fig. 3D**).

T391 is positioned near the active site, closer to the small subunit interface than the C-terminus (**Fig. 4E**). The hydroxyl group of the threonine side chain of residue 391 forms a hydrogen bond with the peptide backbone at M387, stabilizing the α-helix. This interaction is eliminated upon substitution with the non-polar amino acids isoleucine or valine. I251 uniquely faces the internal cavity of the holoenzyme and varies as methionine, valine, or leucine at this position across related enzymes and is one of the most positively selected residues in photosynthetic RbcL (*66*). Both M251 and V251 were not present in the screened library and will require further studies to determine if variation at this site is a catalytic switch.

Substitutions T391I and A438R may alter the active site architecture in a way that reduces the activation energy of CO_2_ addition to RuBP, or increases the activation energy of O_2_ addition (*67*), which may underlie the observed improvement in 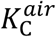. Similarly, T391I may influence structural rearrangements of the active site during catalysis to accelerate CO_2_ addition as an explanation for the marked improvement to 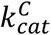 exhibited by this variant. Future detailed kinetic characterization of oxygen affinity (*K*_O_), and the derived 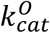, will prove useful in understanding how mutations at position 438 and 391 influence the oxygenation properties of tobacco Rubisco.

### Improved *Nt*RbcL variants can assemble *in planta*

To further position the *Nt*RbcL variants within the context of Rubisco evolution, we conducted a detailed phylogenetic analysis of the amino acid identity at selected sites (figure S7). Only the evolved residues L251 and D468 appear naturally, and only at a low frequency, in vascular plants. In comparison, I391 and V465 are generally observed only in distantly related Rubisco from diatoms and bacteria, respectively, and R438 is virtually never observed despite the catalytic benefit these residues confer to *N. tabacum* Rubisco.

While these substitutions allow Rubisco production in a heterologous bacterial system, we wondered if biogenesis disruption in chloroplasts may explain their absence in plants. Notably, the production of heterologous plant RbcL in leaf chloroplasts is challenged by the protracted timelines to generate a chloroplast genome (plastome)-transformed line. We therefore developed a transgenic *N. benthamiana* line ^*Nb*^*rbc*M that is suited to rapidly screen the chloroplast assembly compatibility of RbcL via *Agrobacterium*-mediated transient expression. This ^*Nb*^*rbc*M line was generated by plastome transformation with plasmid p^cm^trLA, previously used to enable the same replacement in *N. tabacum* cv. Petit Havana (*68*) (**Fig. 5A**). In ^*Nb*^*rbc*M the native *rbc*L plastome gene is replaced with a codon-modified ^cm^*rbc*M gene coding the smaller (100 kDa) *R. rubrum* Form II Rubisco dimer (RbcL_2_) along with the *aad*A gene coding for spectinomycin resistance. Both ^*Nb*^*rbc*M lines generated were spectinomycin-resistant and required elevated CO_2_ for growth in soil, owing to the poor catalytic efficiency and oxygen sensitivity of *R. rubrum* Rubisco (fig. S8).

**Fig. 5.**
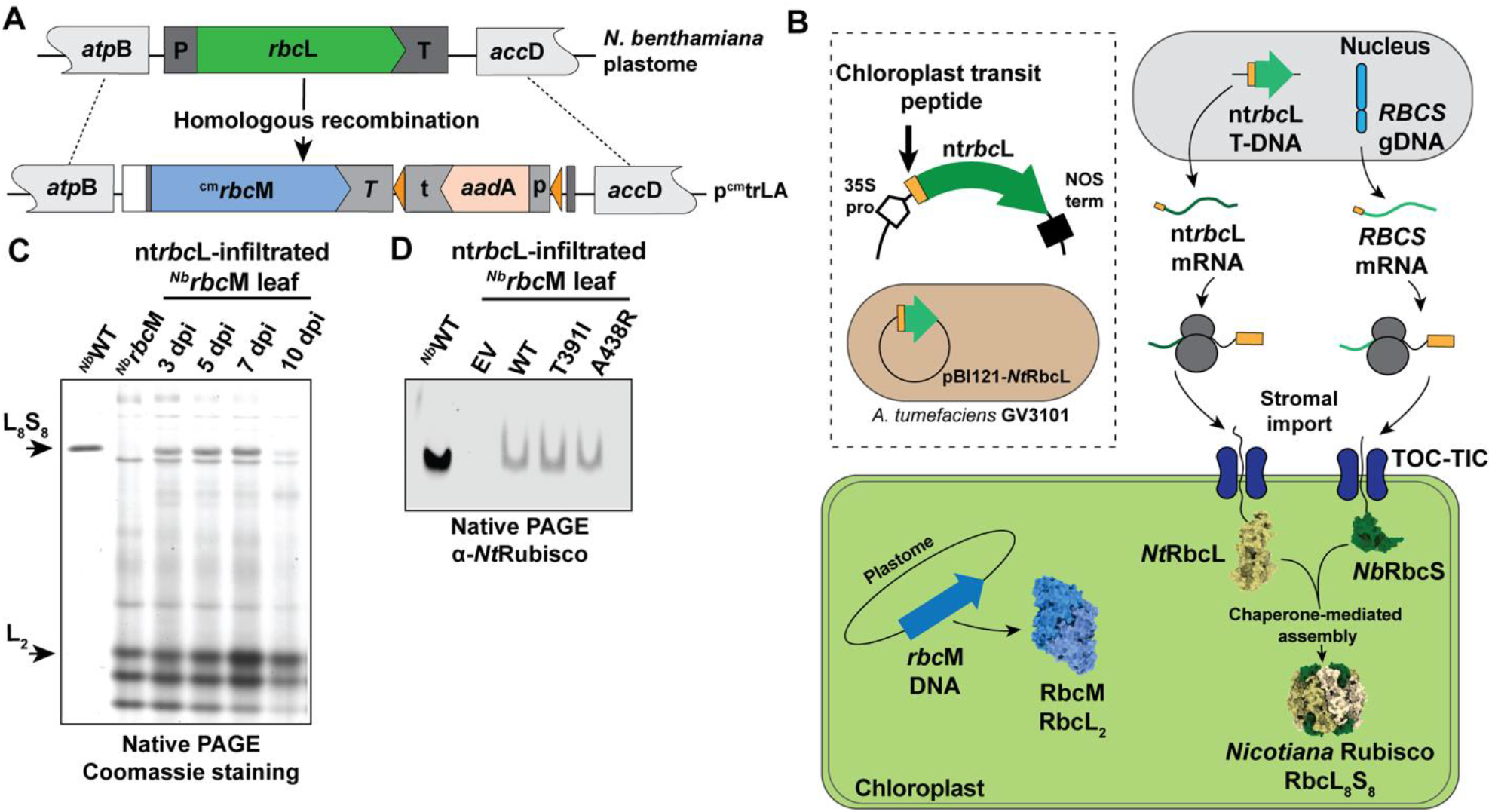
*Nt*RbcL variants fold and assemble in leaf chloroplasts. (**A**) Plastome transformation of *N. benthamiana* with plasmid p^cm^trLA directed the replacement of *rbc*L with a codon-modified *rbc*M (^cm^*rbc*M) gene and *aad*A spectinomycin resistance marker gene to generate independent ^*Nb*^*rbc*M lines (fig. S8). P, 5’ untranslated region (UTR) and first 42 bp of *rbc*L coding sequence; T, 285 bp of 3’ UTR; *T*, 222 bp of *psb*A 3’ UTR; p, 16S rDNA *rrn* promoter and 5’ UTR; t, *rps*16 3’ UTR (*68*). Orange triangles represent loxP sites for marker excision, if desired. (**B**) Schematic of transient RbcL expression and RbcL_8_S_8_ production in the ^*Nb*^*rbc*M chloroplasts via Agroinfiltration of plasmid pBI121-*Nt*RbcL. Shown are how the endogenous nuclear *RBCS* mRNA copies and *rbc*L transgene are translated in the cytosol as precursor peptides coding equivalent N-terminal chloroplast transit peptides that facilitate their import into the chloroplast stroma for assembly into the Rubisco holoenzyme. (**C**) Native PAGE of leaf soluble protein isolated from pBI121-*Nt*RbcL Agroinfiltrated leaves 3–10 days post infiltration (dpi). Infiltrated leaf protein was loaded at 5-fold excess relative to wild-type sampled leaf area to visualize production of ∼520 kDa tobacco L_8_S_8_ Rubisco and the 100 kDa *R. rubrum* L_2_ Rubisco via Coomassie staining. (**D**) Native PAGE Western blot against tobacco Rubisco for native, wild-type *N. benthamiana* (^*Nb*^WT) and ^*Nb*^*rbc*M expressing wild-type and variant *Nt*RbcL. EV; empty vector sampled from an equivalent leaf area.

As seen in equivalent *R. rubrum* Rubisco-producing *N. tabacum* (*68*) and *Solanum tuberosum* (*69*) (potato) genotypes, the ^*Nb*^*rbc*M lines retained RbcS and Rubisco chaperone production from their nuclear-encoded genes (**Fig. 5B**). Agroinfiltration of plasmids coding *N. tabacum* RbcL and the 57 amino acid N-terminal chloroplast-transit peptide from RbcS thus allows for the transient production of RbcL_8_S_8_ complexes within the chloroplasts of ^*Nb*^*rbc*M, as confirmed by native PAGE screens that showed tobacco holoenzyme production between 3–7 days post-infiltration (dpi; **Fig. 5C**).

Infiltration of equivalent constructs coding the I391 and R438 substitutions yielded detectable levels of assembled Rubisco at 5 dpi (**Fig. 5D**). Thus, despite these residues being extremely rare in natural plant Rubisco lineages (table S1 and fig. S7), their production is supported in a chloroplast environment.

Leaf photosynthesis rates in C_3_ plants can be simulated by integrating *in vitro* derived Rubisco catalytic parameters into the established biochemical Farquhar, von Caemmerer, and Berry model (*12, 13, 70-73*). In chloroplasts producing equivalent amounts of Rubisco (20 μmol catalytic sites m^−2^), the *Nt*RbcL^T391I^ mutant is predicted to support the greatest gain in Rubisco activity-limited (*A*_c_) photosynthetic potential. *Nt*RbcL^I465V^ and *Nt*RbcL^N468D^ variants provided the next highest rates after *Nt*RbcL^T391I^ (**Fig. 6**). As the *S*_C/O_ of each variant remained equivalent to wild-type Rubisco (**Table 1**), the electron transport-limited rates of photosynthesis (*A*j; when light-dependent RuBP production becomes most limiting) are predicted by the model to arise at lower chloroplast CO_2_ concentrations (*C*_c_). Nevertheless, each mutant is predicted to support between 15 to 25% faster CO_2_ assimilation rates under current atmospheric CO_2_ levels of 420 ppm, which correspond to *in planta C*_c_ pressures between ∼250–320 µbar.

**Figure 6.**
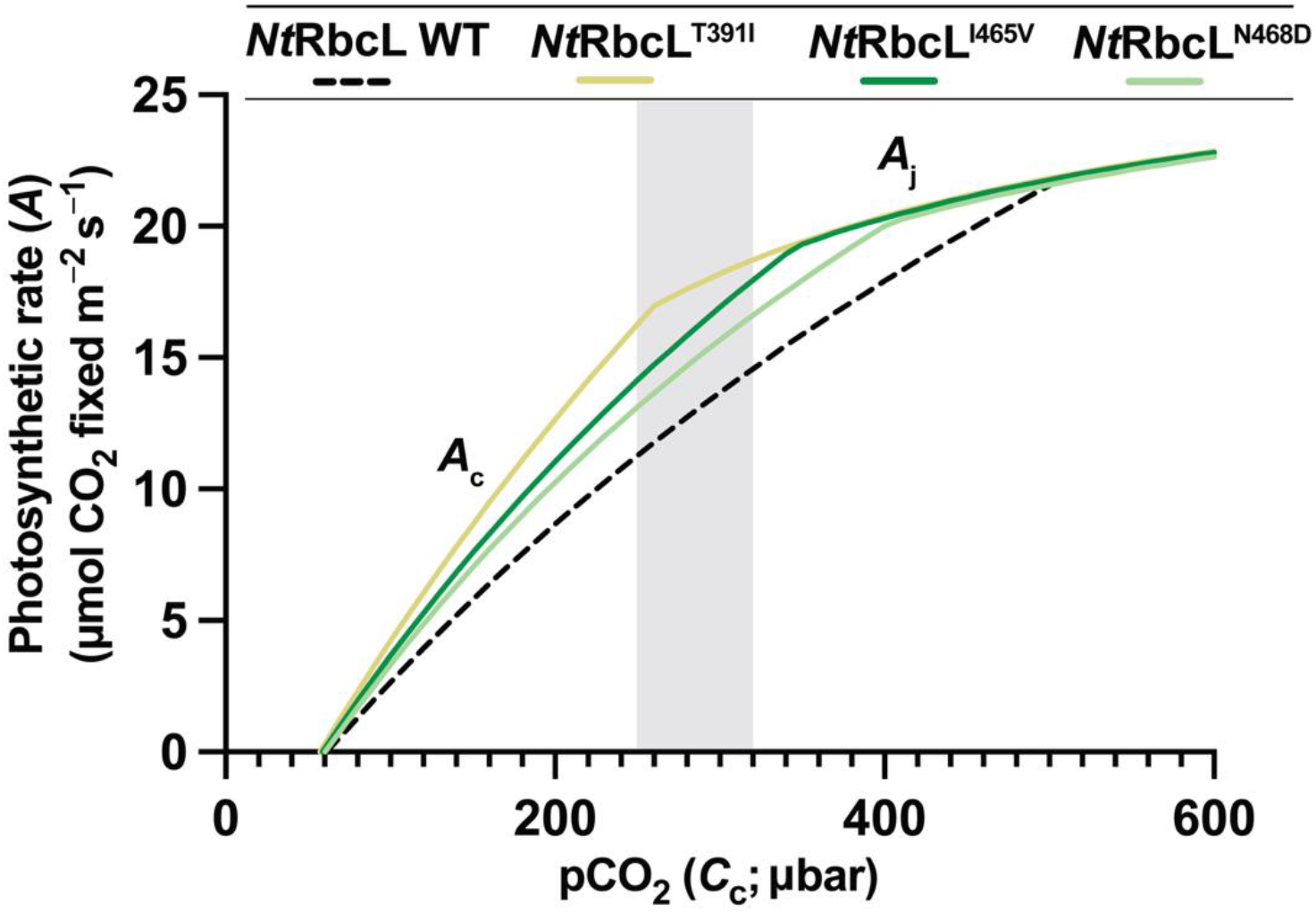
*Nt*RbcL variants support higher modeled rates of photosynthesis in C_3_ leaves. (**A**) Modeled Rubisco activity limited (*A*_c_) and electron transport-limited (*A*_j_) photosynthetic assimilation rates at 25°C in C_3_ leaves expressing variant or wild-type tobacco Rubisco using the kinetic parameters listed in **Table 1**. Photosynthetic simulations are as described by (equations S1 and S2). pCO_2_ (x-axis) represents the partial pressure of dissolved CO_2_ in a C_3_ chloroplast and shaded region represents typical CO_2_ concentration of a C_3_ chloroplast (250–320 µbar) under current atmospheric CO_2_ (420 ppm).

## DISCUSSION

In this work, we used ML to inform the directed evolution of tobacco Rubisco, yielding variants with improved 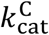 and 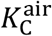 *in vitro*. By using the structure-informed pLM ESM-IF1 to recommend mutagenesis sites for directed evolution libraries, we were able to focus sequence space exploration toward residue changes more likely to be functionally relevant compared to random mutagenesis (**Fig. 3B**), avoiding the numerous deleterious mutations that would have been sampled using error-prone PCR. This approach allowed for substantial minimization of the starting mutational library to reduce the cost and difficulty of generating and maintaining large sequence libraries. In addition, the use of ML-assisted site-saturation mutagenesis allows for amino acid substitutions that require dinucleotide substitutions in the parental codon that would be under sampled using single round error-prone PCR. This approach may explain why A438R (a GCT to CGT codon substitution) was not identified from nt*rbc*L variants generated from random mutagenesis libraries (*12*).

ESM-IF1 was less conservative in mutational predictions compared to the sequence-only ML ensemble ESM-1bv, due to the high level of primary amino acid sequence conservation present in land plant Rubisco (fig. S1). Models, such as ESM-IF1, trained on structural databases potentially avoid such constraint by focusing attention on biophysical principles of the Rubisco structure rather than existing evolutionary trajectories.

While successful at predicting function-enhancing amino acid sites in *Nt*RbcL, ESM-IF1 did not always predict the highest-performing substitution at each site. For example, *Nt*RbcL^T391I^ was not initially predicted. Only T391R was recommended by ESM-IF1, and that substitution was not enriched following selection in RDE2. On the other hand, some substitutions that enhanced catalysis, such as I465V, were directly predicted, as was the substitution M116L, a substitution that improves 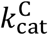 by 30% previously identified during *N. tabacum* Rubisco directed evolution driven by error-prone PCR (*12*). Site-saturation mutagenesis at key ML-recommended sites, in combination with *in vivo* selection, may therefore be a productive strategy for plant Rubisco engineering.

Structure-informed ML through ESM-IF1 suggested substitutions in tobacco Rubisco that are rare in, or absent from, other vascular plant Rubisco sequences (fig. S7). Through ML-assisted directed evolution in *E. coli* we found that tobacco Rubisco enzymes containing these substitutions were not only catalytically enhanced (**Table 1**), but were also assembly-competent in the native chloroplast environment (**Fig. 5D**). Functional substitutions at these sites may be under sampled by methodologies primarily reliant on extant sequence information (*40*), including sequence-only ESM models.

A promising future direction for ML-assisted Rubisco engineering comes from supervised ML models, which input kinetic data alongside structure and sequence information to predict beneficial amino acid substitutions. These models could provide a further advantage when designing Rubisco sequence libraries, as they can also encode known information about key protein–protein interactions *in planta*. Such encoding is essential, as many changes recommended by the unsupervised models we tested, such as R89P and K94E, are known to interrupt interaction with Rubisco activase (Rca) (*74*), (*75*), (*76*), which can cause severe reductions in plant photosynthetic efficiency (*77, 78*). These and other critical protein–protein interactions therefore do not seem to be captured by general unsupervised pLMs, and must be considered when testing substitutions recommended by such models. Supervised models have already been employed to predict extant Rubisco kinetics (*40*), and our results here indicate they may also be useful when used to generate variant predictions or entirely redesigned sequences.

The best-performing variant we isolated, *Nt*RbcL^T391I^, displayed an ∼43% higher carboxylation efficiency in air than wild-type *Nt*RbcL, mostly owing to its faster carboxylation rate. This rate increase was achieved without compromising *S*_C/O_, an increasingly familiar outcome of carefully analyzed Rubisco directed evolution studies (*11-14, 28*). Interestingly, other selected amino acid substitutions at position 391 in *Nt*RbcL (T391C and T391V) did not improve 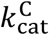, but did benefit the enzyme’s affinity for CO_2_ via reductions in 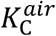 relative to wild-type *Nt*RbcL, suggesting that this locus may be an important modulator of catalysis that is unexplored in plant Rubisco sequence space. Notably, the original substitution predicted by ESM-IF1 (T391R) was not enriched by selection in RDE2. The remaining improved variants displayed enhanced 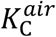 leading to significant improvement to carboxylation efficiency for the I251L, A438R and I465V substitutions.

Large-scale experimental surveys of Rubisco catalysis from vascular plants demonstrate significant variation in each catalytic parameter across natural isoforms (*79*) which encompass the variation exhibited by single-site substitutions characterized in this study. These surveys broadly found *S*_C/O_ to correlate positively with 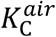 and negatively with carboxylation efficiency in air 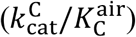 (*43*). However, up to threefold variation in 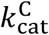 and 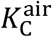 can be observed among enzymes with relatively similar *S*_C/O_ values. The improvements presented in this study, alongside those observed from ancestral reconstructions of Rubisco from the *Solanaceae* family (*22*), show that engineered enhancements are accessible and have the potential to result in improved leaf photosynthetic rates at physiological CO_2_ concentrations.

In conclusion, we used ML-assisted directed evolution to enhance the carboxylation kinetics of tobacco Rubisco through improvements to both carboxylation rate and affinity for CO_2_ in air. It remains possible that factors such as reductions in chloroplast Rubisco content (*80*), increased xylulose-1,5-bisphosphate (XuBP) production (*81*), and reduced activation state (*74, 80*) caused by these substitutions *in planta* may have prevented their evolutionary adoption in plants, especially given their predicted benefits to C_3_ photosynthesis (**Fig. 6**). Construction and full analysis of stable plastome-transformed *N. tabacum* lines expressing these mutant *rbc*L alleles will be necessary to analyze and understand these potential limitations, as well as how the kinetic changes impact plant photosynthesis and productivity.

This work supports continued efforts to engineer plant Rubisco, including further rounds of ML-facilitated evolution of tobacco and other crop Rubisco, and expanding the explored sequence space to include RbcS due to its influential role on substrate specificity (*82*). Ultimately, successful translation of improved Rubisco kinetics *in planta* has the potential to improve crop photosynthesis toward enhancing crop yields to meet rising global food needs.

## MATERIALS AND METHODS

### ESM model implementation

ESM-1bv recommendations were generated as described (*34, 35*), outputting 24 amino acid substitutions. *In silico* DMS using ESM-IF1 was performed as described (*35*), and scores are available in table S1. All relevant code for ESM recommendations and figure generation is available at https://github.com/julielmcdonald/RubiscoMLDE.

### Sequence alignment and conservation analysis

We used the Streptophyta sequence dataset curated by Bouvier *et al*. (*9*) to calculate land plant conservation scores relative to *Nt*RbcL. 69 sequences in this dataset were aligned using MAFFT (*83*), and the conservation score at each residue was calculated as the percentage of sequences in the alignment that contained the same amino acid at a given position relative to *Nt*RbcL.

To calculate the conservation score for all species, as shown in **Fig. 1E**, Jackhmmer (*84*) was used to search for sequences with homology to *Nt*RbcL in the UniProtKB/Swiss-Prot database, yielding 740 representative sequences for conservation analysis. Sequences coding for Rubisco-like proteins were manually removed, resulting in an alignment of 679 protein sequences from bacteria, algae, and plants. Conservation scores were then calculated as above.

To determine which recommendations from ESM were rare in plant Rubisco (table S1) and for alignment of winning sites (fig. S7), RbcL sequences from varying taxonomies were extracted from GenBank and aligned using MAFFT. Full analyses, including code used to survey residues at each locus, are available at https://github.com/julielmcdonald/RubiscoMLDE.

### Variant library generation and selection

For the ESM-IF1 site-saturated variant library, gene fragments containing saturated mutants at selected sites were obtained from Twist Biosciences and were amplified using the primers gg-1 and gg-2. The plasmid pET-*Nt*LS-Rca was amplified using the primers gg-3 and gg-4 (table S7). Fragments were PCR-purified (Omega) and assembled using Golden Gate Assembly via *Bsa*I-HF (New England Biolabs) cutting and T4 ligase (New England Biolabs) ligation overnight. The product of the Golden Gate reaction was purified (Omega), 100 ng was electroporated into NEB^®^ Stable *E. coli* (New England Biolabs), and the outgrowth was plated onto ten Luria broth (LB) agar plates containing 100 µg/mL ampicillin. After 2 d of growth at 30 °C, approximately 400 colonies were scraped from the plates and the plasmid pool was purified (Omega).

For selection of both libraries, 150 ng of library was electroporated into BL21 Star™ *E. coli* (Invitrogen) containing the plasmids pACYC^*prk:nptII*^ and pCDF-*Nt*Asmbl. The cells were grown at 37 °C for 1.5 h before plating on selective agar plates containing 100 µg/mL ampicillin, 100 µg/mL spectinomycin, 35 µg/mL chloramphenicol, 50 µg/mL tetrazolium chloride (TTC), 0.2 mM isopropyl-β-D-thiogalactopyranoside (IPTG), 400 µg/mL kanamycin, and increasing concentrations of L-arabinose (0.05%–0.2% [w/v]). Plates were incubated for 2 d at 25 °C in air supplemented with 0.5% [v/v] CO_2_. All colonies from each selection condition were scraped and the plasmid pool was purified for NGS. Variant nt*rbc*L selected for RDE rescreening were generated using the Q5 site-directed mutagenesis kit (New England Biolabs) or were synthesized by Genscript, Inc.

### Next-generation sequencing

Plasmids from the input library and all selection conditions were linearized, then sequenced and analyzed as described previously (*11*). We used the Genome Analysis Toolkit (GATK) (*85*) to identify single nucleotide polymorphisms (SNPs) in pET-*Nt*LS-Rca. Mutations at 0.10% or higher frequency (determined using a read quality score of 30) in all samples, including the input library, are detailed in table S5.

### RDE spot-testing

Colonies of pET-*Nt*LS-Rca, pCDF-*Nt*Asmbl, and pACYC^*prk:nptII*^ co-transformed into BL21 Star™ *E. coli* (Invitrogen) were grown on LB plates containing 100 µg/mL ampicillin, 100 µg/mL spectinomycin, and 35 µg/mL chloramphenicol (LB-ACS). A colony was used to inoculate 1 mL of LB-ACS, which was then incubated at 37 °C and 180 rpm to an optical density at 600 nm (OD_600_) of ∼0.2. The resulting culture was diluted with LB to an equivalent OD_600_ of 1×10^-4^ and 20 µL was spotted onto an LB-ACS plate containing 50 µg/mL TTC, 0.2 mM IPTG, 400 µg/mL kanamycin, and incremental concentrations (0–0.2% [w/v]) of L-arabinose. The plates were incubated at 25 °C in air supplemented with 0.4% [v/v] CO_2_. Growth was monitored over 8 d.

### Histidine-ubiquitin-tagged Rubisco purification

Calcium-competent BL21 Star™ *E. coli* were co-transformed with pET-*Nt*LS-Rca (containing wild-type or mutated nt*rbc*L), pCDF-*Nt*Asmbl, and pACYC-*Nt* ^His6^Ubq-RbcS. A colony from this transformation was picked and grown in 25 mL of LB-ACS overnight. 20 mL of overnight culture was used to inoculate 1 L LB-ACS. This culture was grown at 37 °C at 180 rpm to OD_600_ = 0.8 (∼2.5 h), at which point protein expression was induced via addition of IPTG to a final concentration of 0.5 mM and the temperature was reduced to 23 °C. Growth continued for 18 h, and then cells were pelleted by centrifugation for 15 min at 4000 × *g* at 4 °C. Cell pellets were snap-frozen in liquid N_2_ for storage at −80 °C.

For Rubisco purification, the stored cell pellets were resuspended in 20 mL ice cold lysis buffer [30 mM EPPS-NaOH at pH 8.0, 150 mM NaCl, 10 mM imidazole (Alfa Aesar), 1 mM phenylmethylsulfonyl fluoride (PMSF; Amersco), 2 mM D,L-dithiothreitol (DTT; Sigma-Aldrich) and 1 µL of DNase (Sigma-Aldrich)], then lysed using a needle-tipped sonicator (30% amplitude with six 20 s on/off cycles) on ice. Following centrifugation (20,000 × *g* for 30 min at 4 °C), the soluble protein was filtered through a 0.22 µm filter before immobilized metal affinity chromatography purification using a 5-mL HisTrap™ excel column (Cytiva) equilibrated with lysis buffer. After loading soluble lysate, the column was washed with 100 mL lysis buffer containing 60 mM imidazole. Rubisco proteins were then eluted with 12 mL lysis buffer containing 300 mM imidazole. For ubiquitin tag cleavage, 400 µL of 80% [v/v] glycerol, 40 µL of 1.5 M β-mercaptoethanol (Sigma-Aldrich), and 150 µL of 2 mg/mL USP2 protease was added to the elution. The resulting solution was incubated at 30 °C for 30 min to initiate cleavage of the ubiquitin tag, then dialyzed at 4 °C overnight against dialysis buffer (10 mM EPPS-NaOH at pH 8.0, 150 mM NaCl, 1 mM EDTA, and 5% [v/v] glycerol). USP2 protease was purified as previously described (*57*).

The dialyzed protein was subjected to IMAC using a 5 mL HisTrap™ excel column equilibrated with dialysis buffer to remove the histidine-tagged USP2 protease and any uncleaved Rubisco. Untagged, pure Rubisco was collected in the flow-through and was concentrated using an Amicon^®^ Ultra Centrifugal Filter (Sigma-Aldrich) with a 100 kDa molecular weight cutoff. Protein was buffer-exchanged into storage buffer (30 mM EPPS-NaOH, pH 8.0, 150 mM NaCl, 1 mM EDTA, 1 mM DTT) and was snap-frozen for future use.

### Untagged Rubisco purification from *E. coli* lysate

BL21 Star™ *E. coli* cultures were grown and induced as described above, with the omission of the plasmid pACYC-*Nt* ^His6^Ubq-RbcS and chloramphenicol in the growth medium. Cells were resuspended in lysis buffer (30 mM Tris-HCl at pH 8.0, 5 mM DTT, 2 mM PMSF, and 1 µL of DNase) and lysis proceeded as above, then lysate was applied to an ENrich™ Q 5 x 50 anion exchange column (Bio-Rad) using a Bio-Rad NGC fast protein liquid chromatography system. Rubisco was eluted from anion exchange using a salt gradient of 0–600 mM NaCl. Rubisco fractions eluted at ∼210 mM NaCl were pooled, concentrated, and loaded onto an ENrich™ SEC 650 10 x 300 column (Bio-Rad) for size exclusion chromatography. Rubisco was eluted using 30 mM Tris-HCl at pH 8.0, 50 mM NaCl, and 10 mM MgCl_2_. Fractions containing Rubisco (fig. S3) were pooled and then snap-frozen for future use.

### Rubisco content and kinetic characterization

Purified Rubisco were used to determine *S*_C/O_ using the [^3^H]-RuBP consumption assay (*86*). ^14^CO_2_-fixation assays to measure 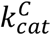 and 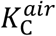 were undertaken in reactions equilibrated with CO_2_-free air at 25 °C containing five incremental ^14^CO_2_ concentrations between 0–300 µM, as previously described (*44, 55*). Briefly, purified *Nt*RbcL protein was diluted to 2 µM in assay buffer (100 mM EPPS-NaOH at pH 8.0, 25 mM MgCl_2_, and 1 mM EDTA) and activated for at least 10 min with 10 mM NaH^14^CO_3_ before diluting 50-fold into a 0.5 mL reaction containing 1.3 µM RuBP. After three min, the reaction was quenched with 50% [v/v] formic acid, unincorporated ^14^C was evaporated overnight, and the fixed ^14^C was counted using a Beckman LS 6500 scintillation counter. Counts were normalized to the number of Rubisco active sites in the purified protein sample, quantified by [^14^C]-CABP binding, as previously described (*55, 87*).

For measurement of Rubisco content in soluble protein of cellular lysates, 200 µL of stationary phase BL21 Star™ *E. coli* co-transformed with pET-*Nt*LS-Rca and pCDF-*Nt*Asmbl was used as a starter culture to inoculate a 10 mL culture of LB-AS in a 50 mL baffled flask. Following growth at 37 °C at 180 rpm to an OD_600_ of 0.8 (∼2.5 h), Rubisco expression was induced with 0.5 mM IPTG and the cells were grown at 23 °C for 18 h before centrifuging for 10 min at 4000 × *g* at 4 °C and snap-freezing the cell pellets in liquid N_2_ for storage at −80 °C. Cell pellets were resuspended in ice cold lysis buffer (30 mM EPPS-NaOH at pH 8.0, 150 mM NaCl, 25 mM MgCl_2_, 10 mM NaHCO_3_, 1 mM PMSF, 2 mM DTT, and 1 µL of DNase) and lysed using a needle-tipped sonicator (20% amplitude with two 20 s on/off cycles) on ice. Following centrifugation (20,000 × *g*, 15 min, 4 °C) the soluble protein was used for ^14^C-CABP binding, as previously described (*55, 87*), and protein quantification via a Pierce™ Bradford Protein Assay Kit against bovine serum albumin. Rubisco concentration as a percentage [w/w] of total cellular soluble protein assumed a molecular mass of 67 kDa for each *Nt*RbcL and RbcS pair. When prepared for kinetics assays, *E. coli* lysates were treated analogously except with the exclusion of 10 mM NaHCO_3_ from the lysis buffer. 20 µL of lysate was activated for at least 10 min with the addition of NaH^14^CO_3_ to 10 mM, then was diluted 1:1 in assay buffer and used for kinetics measurements as described above.

### SDS-PAGE, Western Blotting, and Native PAGE

SDS-PAGE was completed using NuPAGE™ Bis-Tris Mini Protein Gels, 4–12% (Thermo Fisher Scientific), following the manufacturer’s instructions (200 V for 30 min). Native PAGE was completed using Novex™ Value™ Tris-Glycine Mini Protein Gels, 4 to 12% (Thermo Fisher Scientific), following the manufacturer’s instructions (120 V for 2.5 h). Sample loading was normalized to soluble protein content, with 1 µg of protein loaded for Coomassie staining and 0.5 µg of protein loaded for immunoblotting. For immunoblotting, separated proteins were transferred onto a nitrocellulose membrane (Bio-Rad) using a Trans-Blot® Turbo™ Transfer System (Bio-Rad), following the manufacturer’s specifications. Immunoblotting was performed as previously described, using a polyclonal tobacco antibody that reacts with both *Nt*RbcL and *Nt*RbcS (*88*).

### Generation and growth of ^*Nb*^*rbc*M

Replacement of the plastome *rbc*L gene in *N. benthamiana* was achieved via transformation of the plasmid p^cm^trLA (table S6; GenBank AY827488). In p^cm^trLA, 1126 bp and 1176 bp of flanking *N. benthamiana* plastome sequence directs the homologous replacement of *rbc*L and most of its 3’-UTR with the codon-modified *R. rubrum rbc*M gene (^cm^*rbc*M) and an inversely oriented *aad*A selective marker gene (*68*). Pure p^cm^trLA plasmid was coated onto tungsten and transformed into the abaxial side of 10 sterile *N. benthamiana* leaves using a helium accelerator biolistic gun, as previously described (*68*), with 1100 psi rupture discs. After 2 d, the leaves were dissected into ∼1 cm^2^ disks and embedded into selective 1× Murashige and Skoog (MS)/phytagel media containing 0.5 µg/mL spectinomycin, 1% [w/v] sucrose, 1 µg/mL thiamine (Sigma-Aldrich), 0.1 µg/mL α-napthaleneacetic acid (Sigma-Aldrich), and 1 µg/mL 6-benzyl aminopurine (Sigma-Aldrich). The leaf sections were incubated at 25 °C in air containing 2.5 % [v/v] CO_2_.

Three spectinomycin-resistant green calli were dissected after 7 weeks and then transferred to plates of fresh selective MS/phytagel media. The tissue was regenerated for 12 weeks, and then ∼0.1 cm^2^ tissue samples were regenerated again on selective MS/phytagel media. After 17 weeks, the stems of ∼1 cm high plants were cut diagonally, and the plants were propagated into 0.8 L polypropylene pots containing 0.15 L MS/phytagel media supplemented with 0.05 % [w/v] sucrose and 0.25 mg/mL spectinomycin at 25 °C in 2.5 % [v/v] CO_2_-enriched air. 4–6 weeks later, following root development, the plants were transferred to plastic containers filled with a 2:1 dry volume ratio mix of vermiculite/seed-raising soil mix and a teaspoon of osmocote, and were covered with 0.8 L polypropylene pots for 6 d to maintain humidity. After two weeks, the plants were transferred to 2 L pots of soil and were grown to reproductive maturity in a growth chamber at 25/20 °C (16 h light/8 h dark), 150−450 µmol photons m^-2^ s^−1^ illumination, and 2% [v/v] CO_2_-enriched air. Progress towards homoplasmicity was monitored by native PAGE analysis.

### Agroinfiltration of ^*Nb*^*rbc*M

Prior to germination, seeds were frozen at −80 °C overnight, and then were sterilized by incubation in 1% [v/v] bleach for 10 min. Seeds were washed five times with ultrapure water, and then were spread onto 1× MS/phytagel media for germination. Seeds were incubated at 25 °C in a growth chamber equilibrated with air supplemented with 2.0% [v/v] CO_2_ and 25–100 µmol photons m^-2^ s^−1^ illumination for approximately two weeks. Seedlings were then transferred to soil and were grown for approximately three more weeks until suitable for infiltration, with watering approximately once per week.

*A. tumefaciens* GV3101 was used for transient expression of *Nt*RbcL in ^*Nb*^*rbc*M. Plasmids pBI121-*Nt*RbcL and pBI-p19, (containing the gene silencing suppressor P19) were individually transformed into electrocompetent *A. tumefaciens*. Colonies from these strains were grown in LB containing 100 μg/mL kanamycin and 50 μg/mL rifampicin to an OD_600_ of 0.6 (∼24 h). 1 mL of each pBI121-*Nt*RbcL-containing culture was added to 0.5 mL of pBI-p19-containing culture and then pelleted by centrifugation at 15,000 × *g* for 1 min at 24 °C. Cells were resuspended in 3 mL of infiltration buffer (10 mM MES-NaOH at pH 5.5, 10 mM MgSO_4_), and were injected into the leaves of ∼5 week old ^*Nb*^*rbc*M plants using a syringe. Plants were grown for 6 d after infiltration with watering on day 3, and then 0.5 cm^2^ leaf disks were punched and snap-frozen for future use. Leaf samples removed from −80 °C were immediately resuspended in 1 mL of lysis buffer [30 mM EPPS-NaOH at pH 8.0, 150 mM NaCl, 25 mM MgCl_2_, 1 mM PMSF, 2 mM DTT, 1 µL of DNase, and 1% [w/v] polyvinylpolypyrrolidone (PVPP; Sigma-Aldrich)] and were mechanically lysed. After centrifugation at 20,000 × *g* for 15 min at 4 °C to remove insoluble debris, the supernatant was prepared for SDS-PAGE analysis as described above.

### Modeling of photosynthetic rates

Photosynthetic *A*-*C*_c_ curves for C_3_-plants were modeled according to Farquhar *et al*. (*59*) using *in vitro* measures of Rubisco *S*_C/O_, 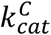 and 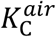 (**Table 1**) (*1, 59*), a leaf Rubisco content of 20 μmol RbcL sites m^−2^, a CO_2_ solubility constant of 0.0334 M bar^−1^, an electron transport rate (*J*) of 120 μmol m^−2^ s^−1^, a non-photorespiratory CO_2_ release rate (*R*_*d*_) of 1 μmol m^−2^ s^−1^, and a stromal O_2_ concentration of 253 μM.

## Acknowledgments

We extend our gratitude to the MIT BioMicroCenter staff for their assistance with high-throughput sequencing experiments, particularly Dr. Vincent Butty, who developed the initial NGS processing scripts.

## Author contributions

J.L.M., B.D.B., S.M.W., M.D.S., and R.H.W. conceived this work. All authors designed experiments and analyzed data or contributed reagents. J.L.M., J.L., Y.Z., B.L.H., R.B, S.M.W., and R.H.W. performed experiments. J.L.M., M.D.S., and R.H.W. wrote the manuscript. All authors edited the manuscript.

## Data, Code, and Materials Availability

All data needed to evaluate the conclusions in the paper are present in the paper and/or the Supplementary Materials. All protein structure visualizations were generated using ChimeraX (*89*); https://www.cgl.ucsf.edu/chimerax/. Raw sequencing reads have been deposited in the NCBI Sequence Read Archive under BioProject accession PRJNA1476684 (https://www.ncbi.nlm.nih.gov/sra/PRJNA1476684). All code is available on Zenodo (DOI:10.5281/zenodo.21269161), and on GitHub (https://github.com/julielmcdonald/RubiscoMLDE). The plasmids pET-*Nt*LS-Rca, pCDF-*Nt*Asmbl, pACYC-*Nt*^His6^Ubq-RbcS, and pACYC^*prk:nptII*^ are available from Addgene (https://www.addgene.org/browse/article/28273150/) with identification numbers 253864, 253865, 253866, and 239580, respectively. Requests for *Nb*rbcM plant lines should be submitted to.

## SUPPLEMENTARY MATERIAL

**Other supporting materials for this manuscript include the following:**

Tables S1, S4, S5 (as .xls files)

## SUPPLEMENTARY TABLES

**Table S1**. ESM-1bv *Nt*RbcL amino acid substitutions, ESM-IF1 *Nt*RbcL amino acid substitutions, and conservation scores across the *Nt*RbcL sequence (see Excel file).

**Table S2.**
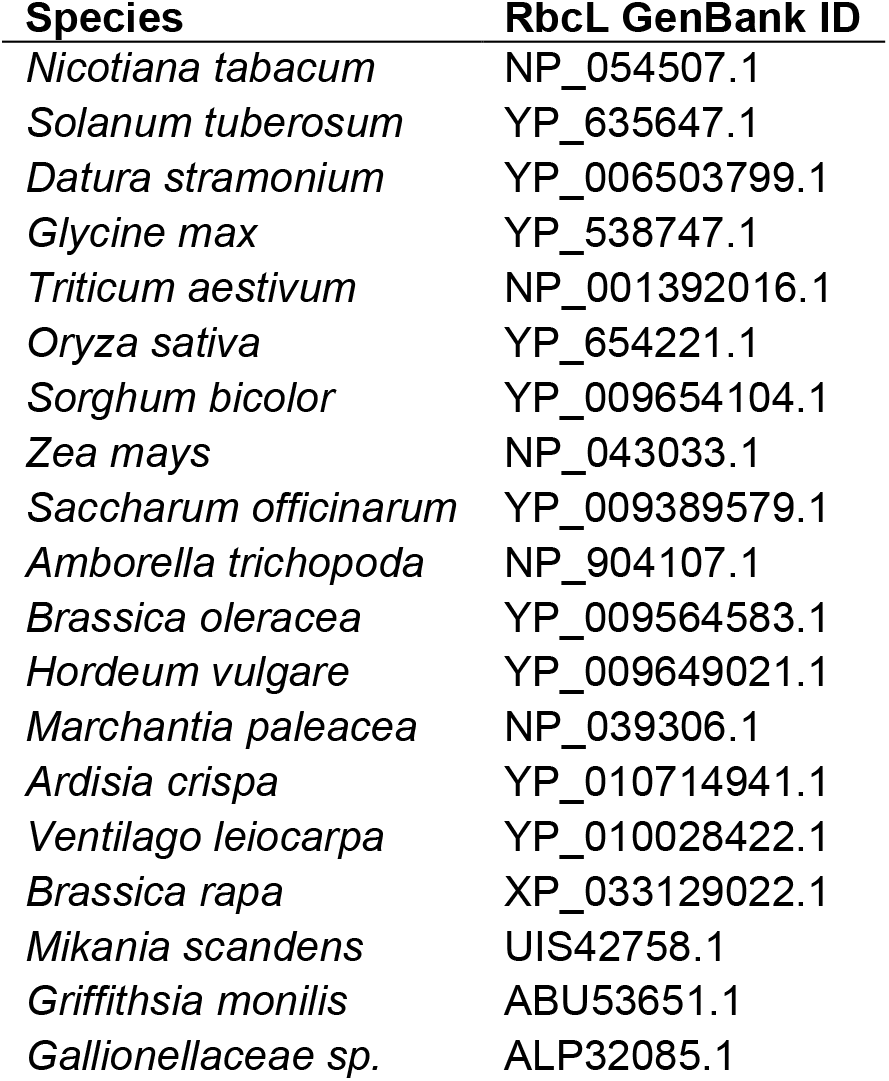
Select RbcL sequences used for alignment.

**Table S3.** Solubility and kinetic characterization of *Nt*RbcL^A369V^. Values provided are means ± standard deviation. N ≥ 3, replicate data is provided in **Supplementary Table 4**. Statistical significance relative to wild-type was calculated via a two-tailed, heteroscedastic *t*-test. ***, *p* ≤ .0005. % TSP; % total soluble protein.

| *NtRbcL* | % TSP | $k_{cat}^C$ (s<sup>-1</sup>) | $K_C^{air}$ (μM) | $k_{cat}^C/K_C^{air}$ (mM<sup>-1</sup> s<sup>-1</sup>) | S<sub>C/O</sub> |
| --- | --- | --- | --- | --- | --- |
| A369 | 1.57 ± 0.7 | 3.50 ± 0.5 | 30.2 ± 4.1 | 117.4 ± 20.9 | 79.7 ± 4.0 |
| V369 | 0.23 ± .1*** | 3.49 ± 1.0 | 25.3 ± 5.2 | 137.2 ± 15.4 | 82.0 ± 4.2 |

**Table S4**. Kinetic characterization of Rubisco variants (see Excel file).

**Table S5**. Characterization of input plasmid library and selected variant frequencies (see Excel file).

**Table S6.** Plasmids used in this study.

| Plasmid | Proteins encoded (GenBank ID) | Source |
| --- | --- | --- |
| pET-*NtLS-Rca* | *NtRbcL* (P00876.2), *NtRbcS* (P69249.1), *NtRca* (Q40460.1) | (24) |
| pCDF-*NtAsmbl* | *NtRaf1* (XP_016475347.1), *NtRaf2* (XP_016516208.1),<br>*NtRbcX* (XP_016509860.1), *NtBSD2* (AYR18863.1),<br>*NtCpn60α* (XP_016509320.1), *NtCpn60β*<br>(XP_016490465.2), *NtCpn20* (XP_016432480.1) | (24) |
| pACYC<sup>prk:nptII</sup> | PRK (WP_011242879.1), NptII (WP_000572405.1) | (14) |
| pACYC-*Nt*<sup>His6</sup>Ubq-*RbcS* | <sup>His6</sup>Ubq (AAV27297.1), *NtRbcS* (P69249.1) | This work |
| p<sup>cmtr</sup>LA | *RbcM* (AAV98336.1), *AadA* (AAV98337.1) | (88) |
| pBI121-*NtRbcL* | *tpNtRbcL* (P00876.2) | This work |
| pBI-p19 | P19 (NP_062901.1) | This work |

**Table S7.**
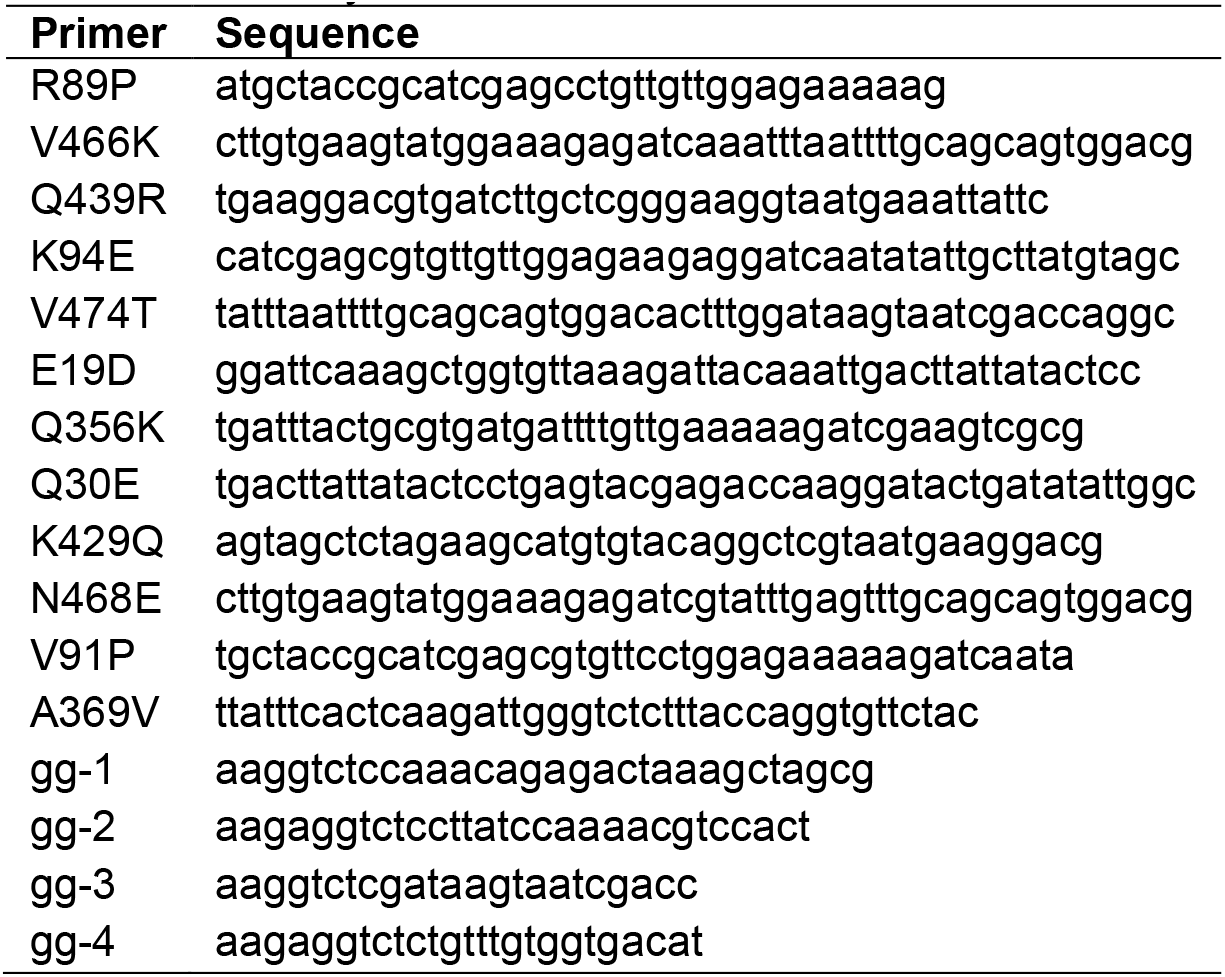
Primers used in this study.

## SUPPLEMENTARY EQUATIONS

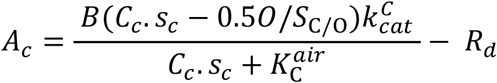

**Equation S1**. *A*_c_; Rubisco activity-limited photosynthesis rate (*1, 59, 90*). *B*; Rubisco catalytic sites per leaf area (20 μmol sites m^−2^). *C*_*c*_; mesophyll chloroplast CO_2_ partial pressure (µbar). *s*_*c*_; CO_2_ solubility in water (0.0334 M.bar^−1^ equivalent to 0.0334 µM.µbar^−1^). *O*; dissolved stromal O_2_ concentration (253 µM). *S*_C/O_; Rubisco specificity factor. 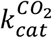; carboxylation rate (s^−1^). 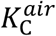; (µM) the apparent Michaelis constant for CO_2_ at ambient (21% [v/v]) O_2_. *R*_*d*_; non-photorespiratory CO_2_ production rate (1 µM.m^−2^.s^−1^). 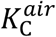 (i.e.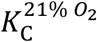) is equivalent to *K*_C_(1+*O*/*K*_O_), as used previously

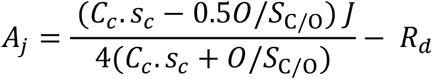

**Equation S2**. *A*_j_, electron transport-limited photosynthesis rate (*1, 59, 90*). Variables described in

**Supplementary Equation 1**. *J*; electron transport rate (120 µmol.m^-2^.s^-1^).

## SUPPLEMENTARY FIGURES

**Figure S1.**
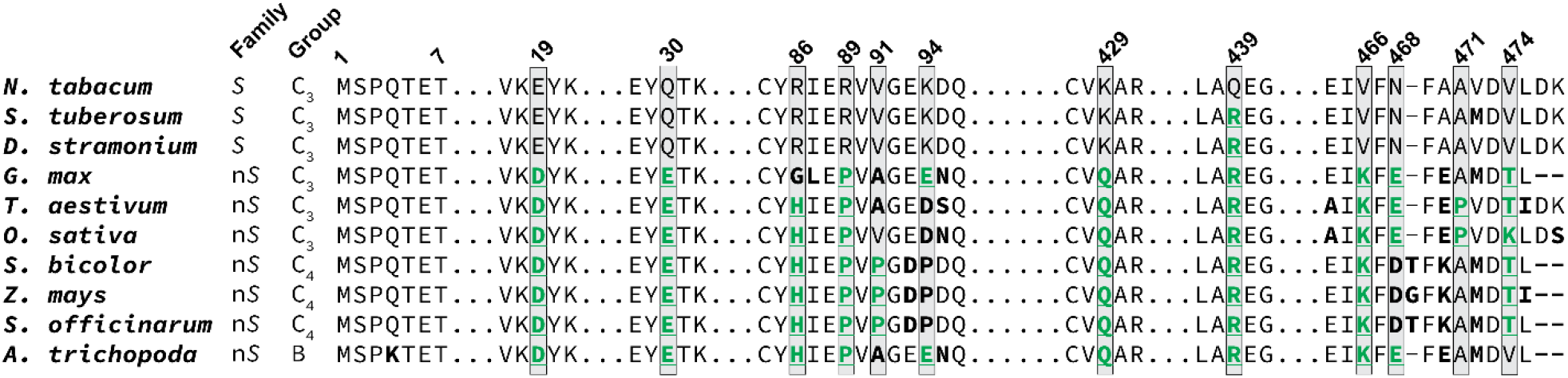
Alignment of tobacco and other plant RbcL sequences. Sites of ESM-1bv recommended variants are boxed and recommended amino acid changes are underlined in green. Sequences were aligned using MAFFT(*83*) under default settings. All sequence differences from tobacco Rubisco are bolded. *S*; *Solanaceae*, n*S*; non-*Solanaceae*, B; basal. Full genus names and GenBank Accession numbers are listed in **Supplementary Table 2**.

**Figure S2.**
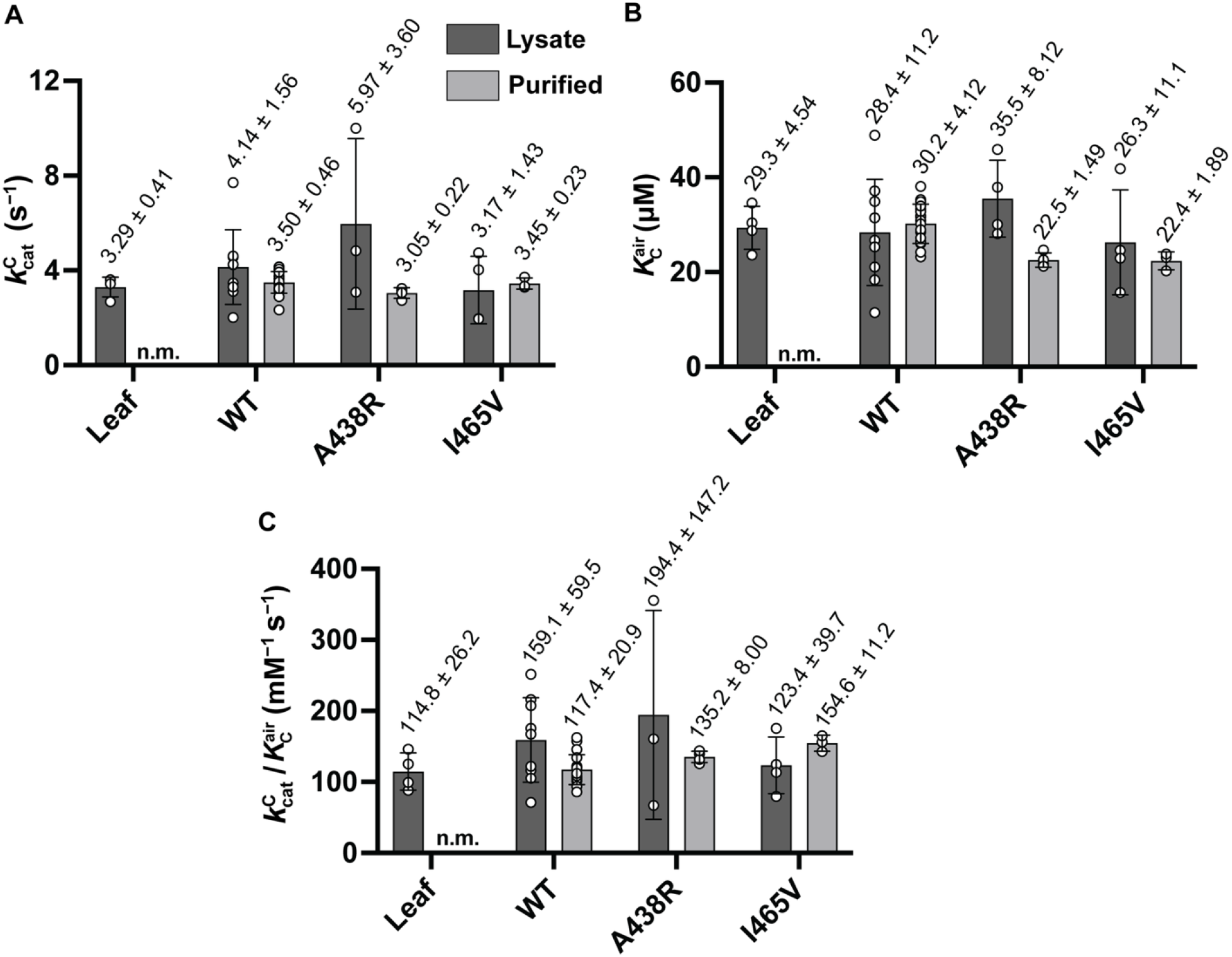
Kinetic measurements for wild-type *Nt*RbcL, *Nt*RbcL^A438R^, and *Nt*RbcL^I465V^ from *E. coli* lysate and purified protein. **A**. Carboxylation rate; 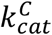. **B**. Affinity for CO_2_ in air; 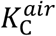 **C**. Carboxylation efficiency in air;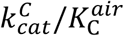. Rubisco from *N. tabacum* leaf lysate was included as a positive control in **A**–**C**. Values shown above bars represent mean ± standard deviation. All differences in kinetics between lysate and purified samples had *p*-values > 0.05 when analyzed via two-tailed, heteroscedastic *t*-test.

**Figure S3.**
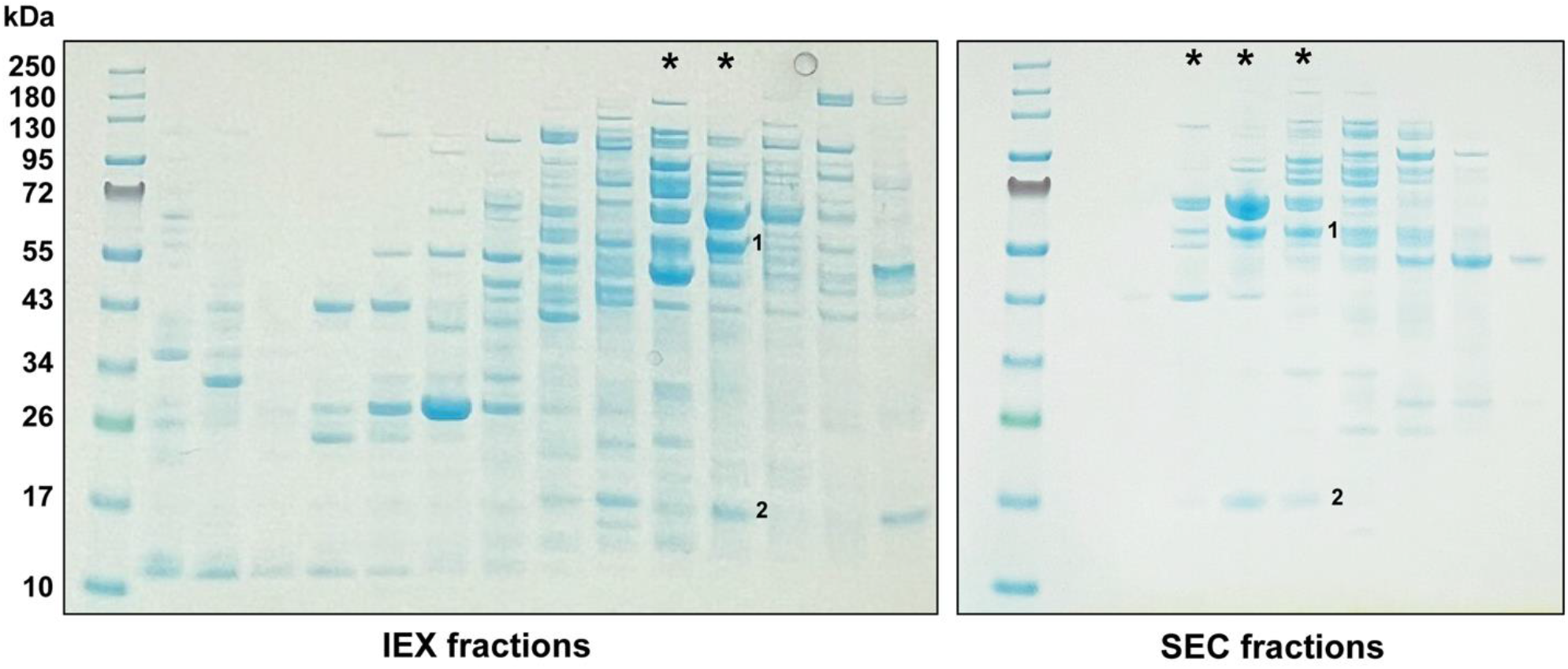
Purification of untagged tobacco Rubisco from *E. coli* lysate. Coomassie-stained SDS-PAGE gels show fractions collected after ion exchange (IEX) and size exclusion chromatography (SEC). Fractions containing Rubisco protein are indicated with asterisks. *Nt*RbcL (1) has a molecular weight of 53 kDa and *Nt*RbcS (2) has a molecular weight of 15 kDa.

**Figure S4.**
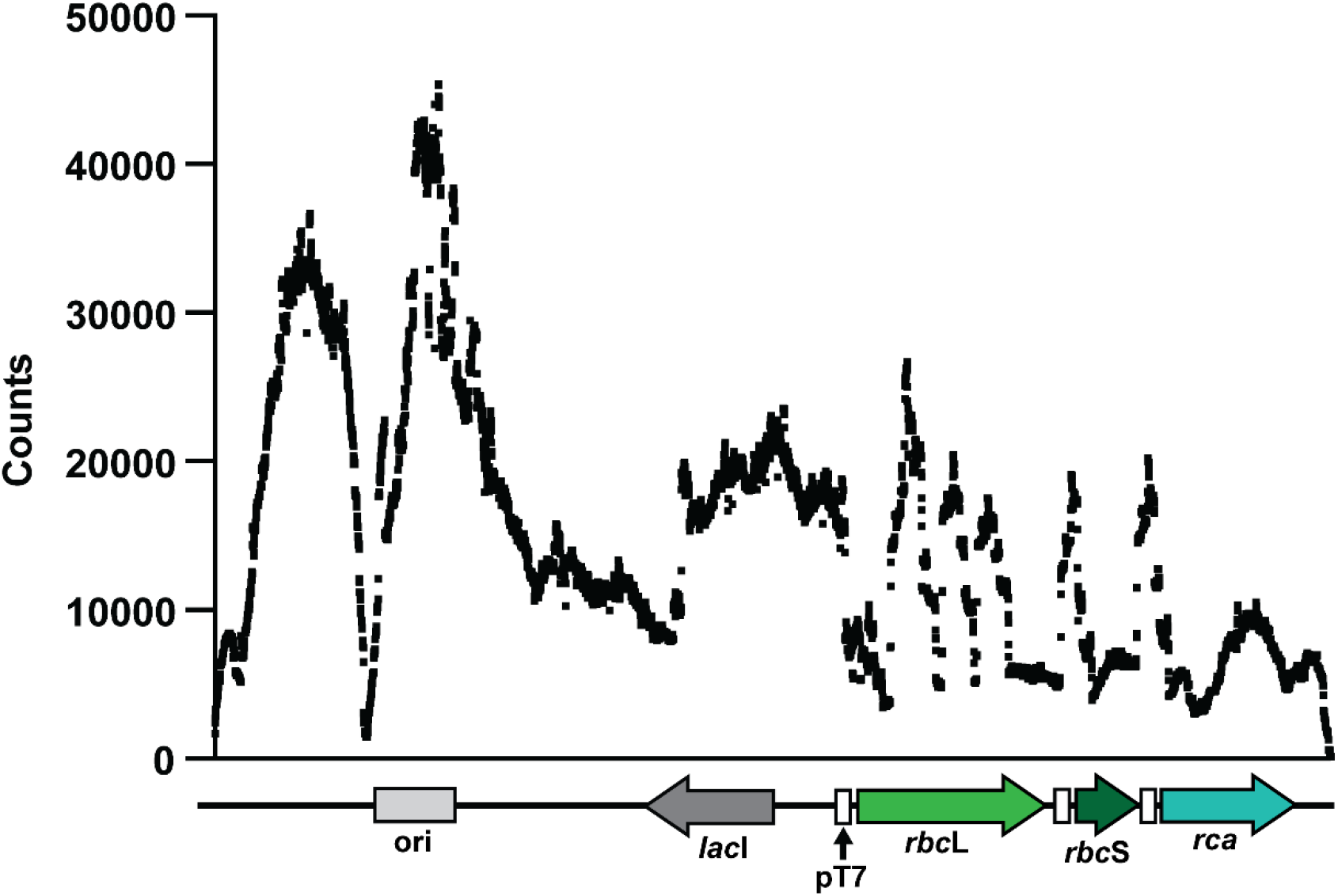
Histidine-ubiquitin-tagged purification of *N. tabacum* Rubisco complexes. **A**. Three-plasmid system for tagged Rubisco expression. Rubisco and assembly plasmids are from Buck *et al* (*24*). **B**. Hypothetical structure of ^His6^Ubq-tagged Rubisco. Rubisco holoenzymes are fused to ^His6^Ubq at the N-terminus of RbcS, enabling pull-down via nickel immobilized metal affinity chromatography (IMAC). Ubiquitin (PDBID: 3U30 (*91*)) was docked onto the tobacco Rubisco structure (PDBID: 4RUB (*50*)) using ChimeraX (*89*). **C**. Purification of wild-type tobacco Rubisco via histidine-ubiquitin tag.

**Figure S5.**
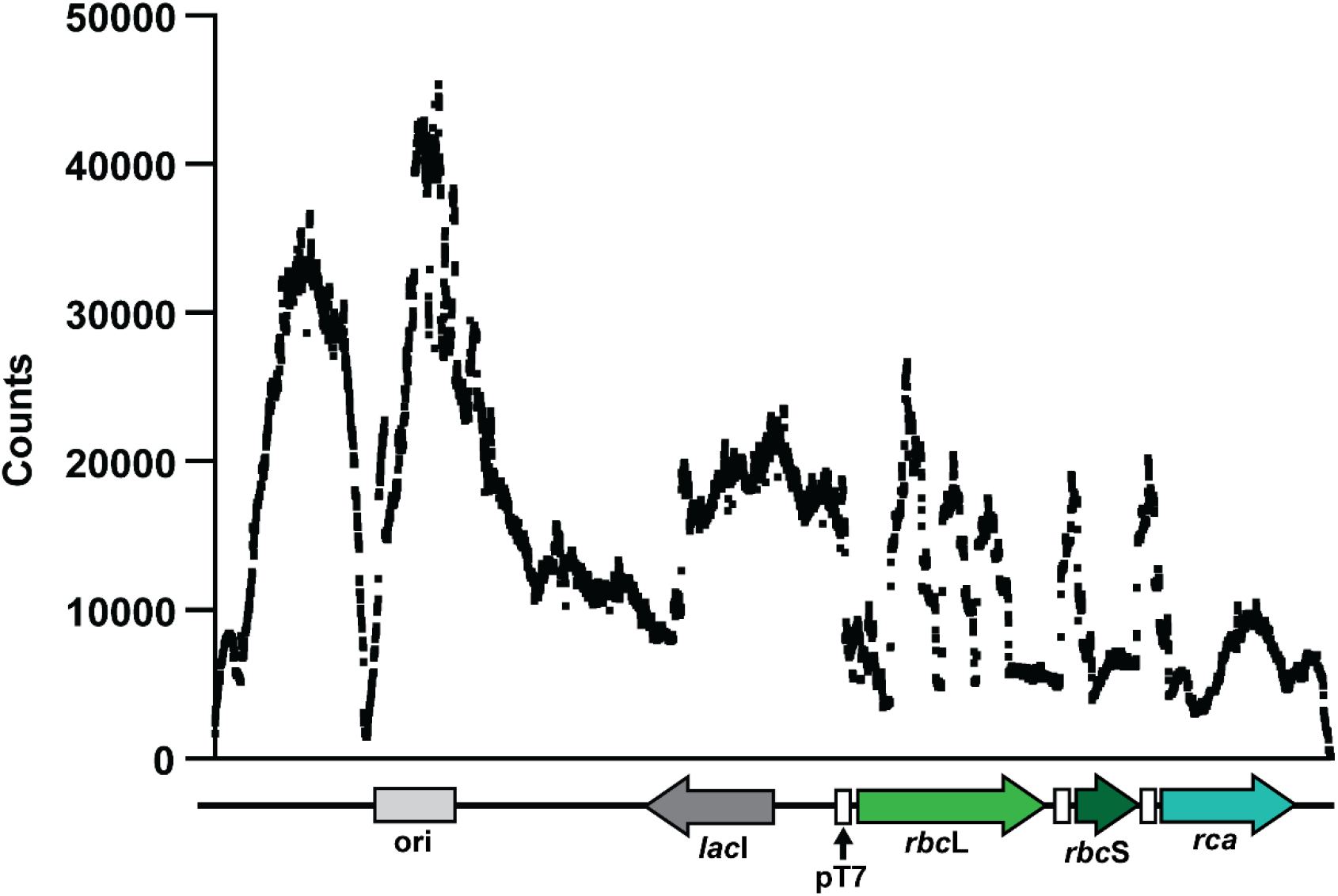
Sequencing depth across pET-*Nt*LS-Rca. Data represents the sum of six total samples: technical duplicates of three independent library preparations. Short-read DNA sequencing was performed using a 150 paired-end kit on an Element Biosciences AVITI™ sequencer, as previously described (*11*).

**Figure S6.**
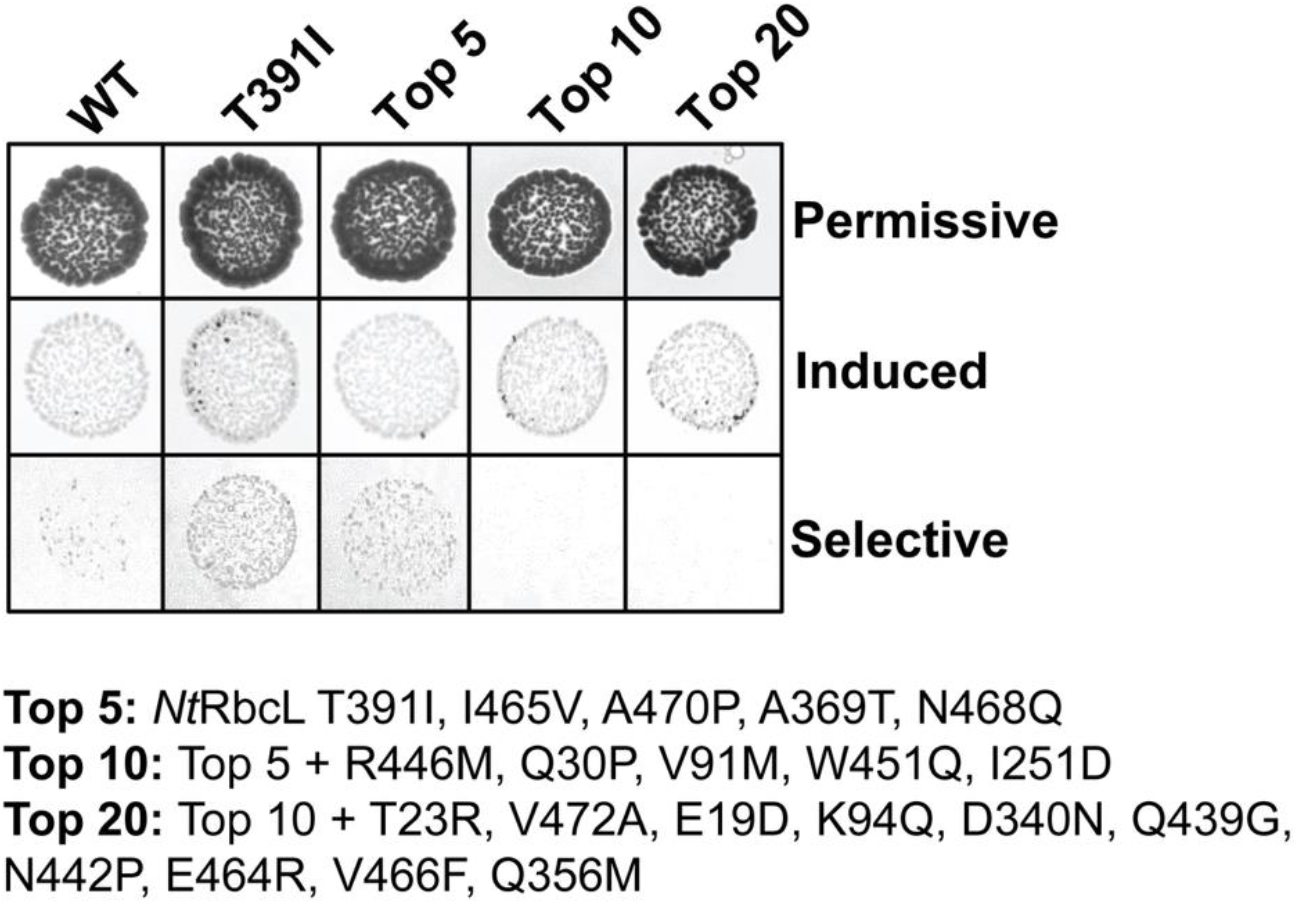
Growth of combinatorial *Nt*RbcL variants in RDE2. Cells were spot-tested on permissive media containing no additives, induced media containing 0.2 mM IPTG for Rubisco expression, and selective media containing 0.15% [w/v] L-arabinose, 400 µg/mL kanamycin and 0.2 mM IPTG for inducing both PRK and Rubisco expression.

**Figure S7.**
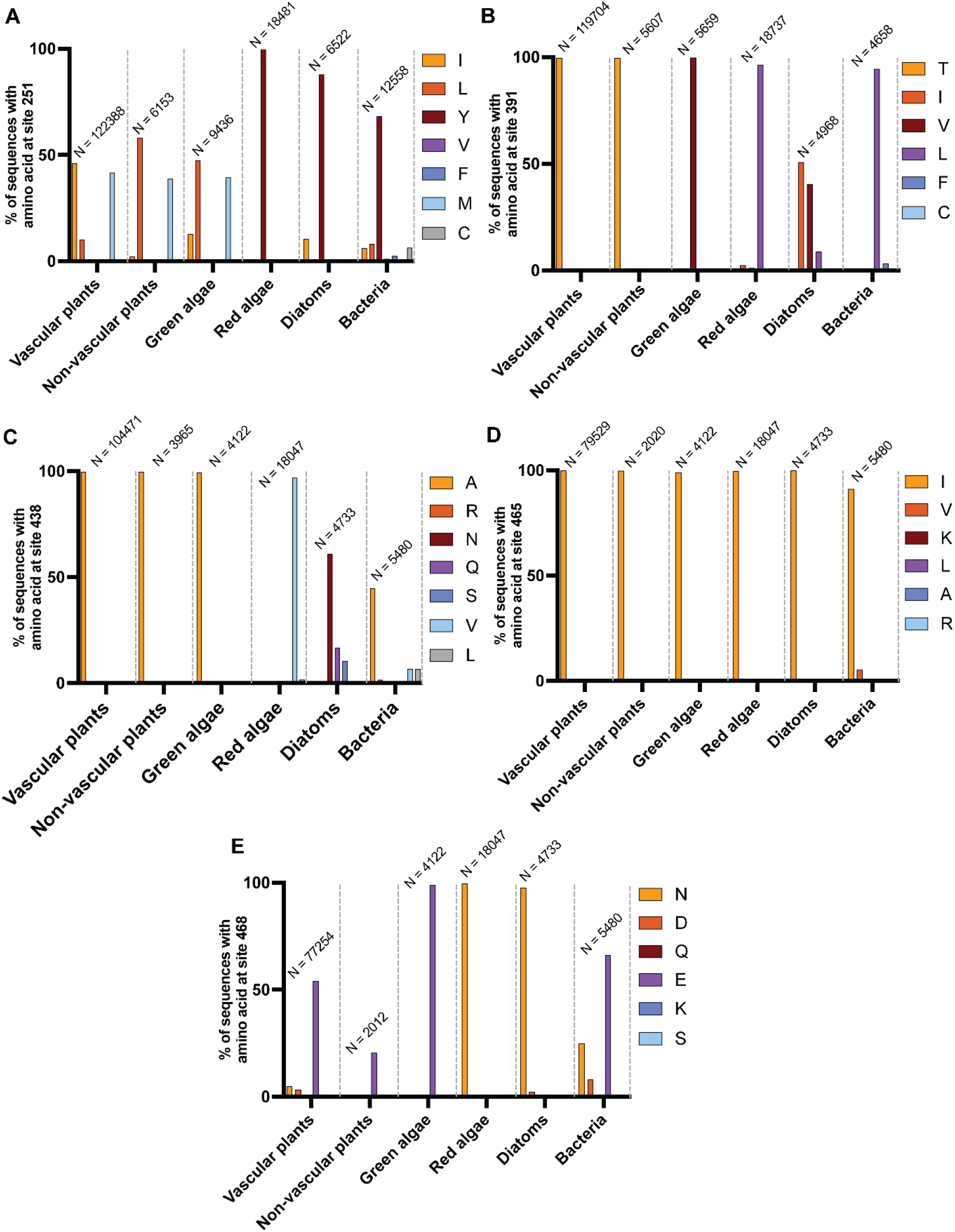
Phylogenetic analysis at position of selected residues. Data are shown as a percentage of the total sequences containing the indicated residue at that position. *N* is equal to the number of sequences that had any amino acid residue at the position; thus partial sequences with missing termini are not represented. **A**. Site 251. **B**. Site 391. **C**. Site 438. **D**. Site 465. **E**. Site 468.

**Figure S8.**
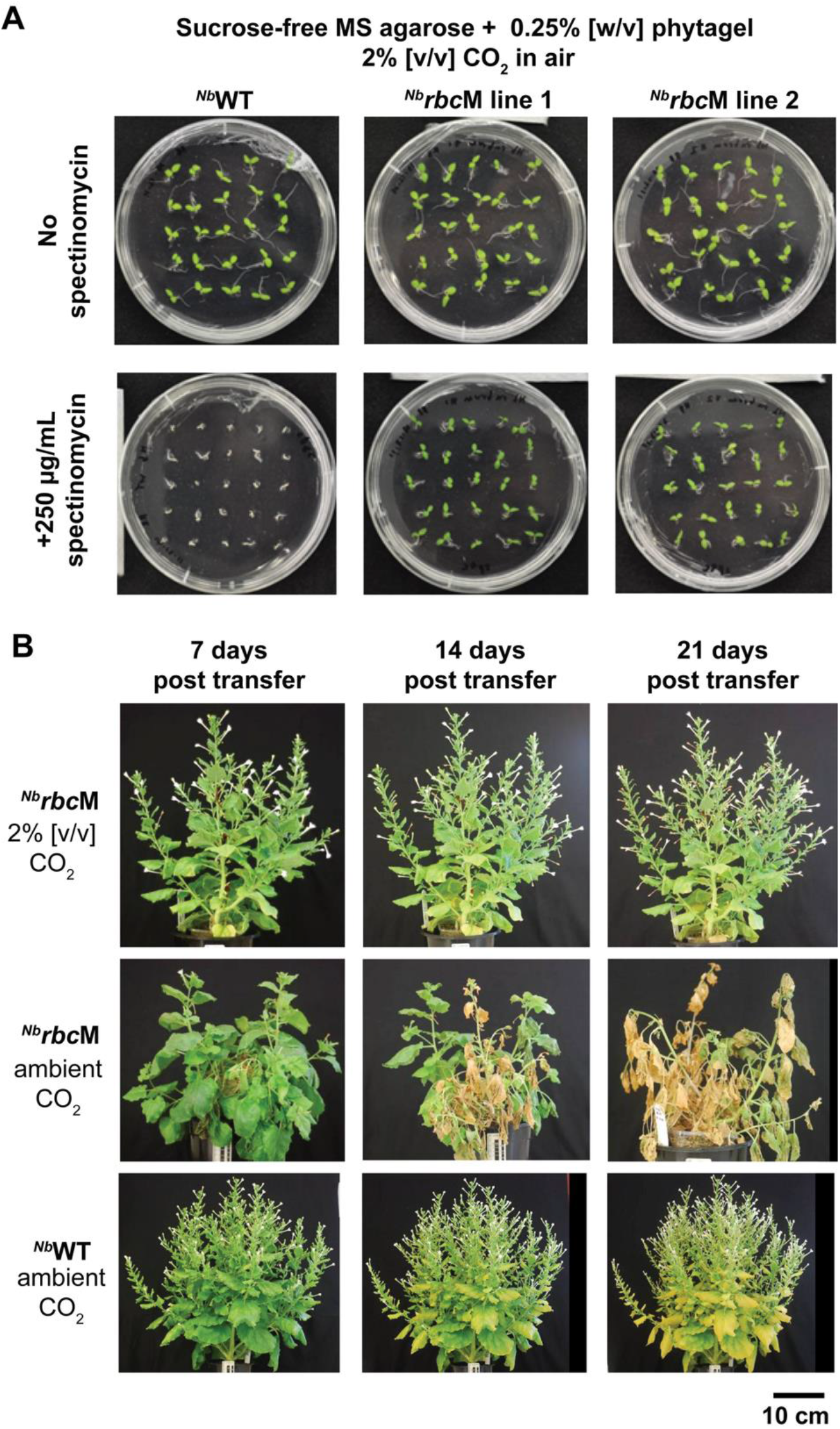
Germination and growth of ^*Nb*^*rbc*M under elevated CO_2_. **A**. Germination of *N. benthamiana* wild-type (^*Nb*^WT) and T_1_ ^*Nb*^*rbc*M plants on media containing spectinomycin shows homoplastomic ^cm^*rbc*M-*aad*A integration. **B**. The T_1_ plants were propagated in soil for 32 days under elevated CO_2_ (air + 2% [v/v] CO_2_) and then either maintained under elevated CO_2_ or grown with identical chamber conditions under ambient CO_2_ (air + 0.06% [v/v] CO_2_) for 7–21 days and imaged. Vegetative growth and seed setting by the ^*Nb*^*rbc*M lines was supported under elevated CO_2_, but not ambient.

